# The structure, binding, and function of a Notch transcription complex involving RBPJ and the epigenetic reader protein L3MBTL3

**DOI:** 10.1101/2022.02.09.479311

**Authors:** Daniel Hall, Benedetto Daniele Giaimo, Sung-Soo Park, Wiebke Hemmer, Tobias Friedrich, Francesca Ferrante, Marek Bartkuhn, Zhenyu Yuan, Franz Oswald, Tilman Borggrefe, Jean-François Rual, Rhett A. Kovall

## Abstract

The highly conserved Notch pathway transmits signals between neighboring cells to elicit distinct downstream transcriptional programs. In given contexts, Notch is a major regulator of cell fate specification, proliferation, and apoptosis, such that aberrant Notch signaling leads to a pleiotropy of human diseases, including developmental disorders and cancers. The canonical pathway signals through the transcription factor CSL (RBPJ in mammals), which forms a transcriptional activation complex with the intracellular domain of the Notch receptor and the coactivator Mastermind. CSL can also function as a transcriptional repressor by forming complexes with one of several different corepressor proteins, such as FHL1 or SHARP in mammals and Hairless in *Drosophila*. Recently, we identified the malignant brain tumor (MBT) family member L3MBTL3 as a bona fide RBPJ binding corepressor that recruits the repressive lysine demethylase LSD1/KDM1A to Notch target genes. Here we define the RBPJ-interacting domain (RBP-ID) of L3MBTL3 and report the 2.06 Å crystal structure of the complex formed between RBPJ, the RBP-ID of L3MBTL3 and DNA. The structure reveals the molecular interactions underlying L3MBTL3 complexation with RBPJ, which we comprehensively analyze with a series of L3MBTL3 and RBPJ mutations that span the binding interface. Compared to other RBPJ-binding proteins, we find that L3MBTL3 interacts with RBPJ via an unusual binding motif, which is sensitive to mutations throughout its RBPJ-interacting region. We also show that these disruptive mutations affect RBPJ and L3MBTL3 function in cells, providing further insights into Notch mediated transcriptional regulation.

## Introduction

Notch is a conserved signaling pathway that is critical for proper metazoan development and homeostasis throughout life^1^. Notch signaling is a transcriptional regulation mechanism whose gene targets regulate diverse cellular processes, such as proliferation, differentiation, and apoptosis, depending on the cellular context of the signal^2^. The pathway is tightly regulated and very sensitive to gene dosage, whereby too much or too little signaling leads to devastating health outcomes. Many human diseases, such as certain forms of congenital syndromes, cancers, and cardiovascular disease, have been linked to mutations in Notch signaling components^1^, and therapeutic modulation of the pathway is an active area of research due to the current lack of long-term solutions^3^. One of the goals of Notch targeted therapeutics is to identify small molecules or biologics that can discriminate between different Notch regulatory transcriptional complexes^4^.

The Notch pathway is activated when a transmembrane Notch receptor on a signal-receiving cell engages with a transmembrane ligand on an adjacent signal-sending cell (Fig 1A)^5^. In mammals there are four Notch receptors (NOTCH1/2/3/4) and five ligands of the DSL (Delta/Serrate/Lag-2) family: JAG1/2 (Jagged1/2) and DLL1/3/4 (Delta-like 1/3/4)^1^. Notch receptors and DSL ligands are large modular multidomain proteins with a single transmembrane spanning region^5^. Ligand-receptor binding triggers endocytosis of the extracellular complex by the signal-sending cell, which exerts a pulling force on the receptor, exposing a cleavage site for ADAM10 (A Disintegrin and Metalloproteinase 10)^6^. ADAM10 cleavage sheds the extracellular domain while the membrane-bound Notch intracellular domain (NICD) is cleaved within its transmembrane region by the gamma-secretase complex, releasing NICD from the cell membrane^5^. Subsequently, NICD localizes to the nucleus where it forms a transcriptional activation complex with the transcription factor CSL (CBF1/RBPJ, Su(H), Lag-1) and a member of the Mastermind family of transcriptional coactivators (MAML1-3, in mammals) (Fig 1A)^7^. The NTC (Notch transcription complex) recruits the mediator complex and the histone acetyltransferases P300/CBP (CREB binding protein) to DNA regulatory elements of Notch target genes to turn “on” transcription^8,9^.

**Figure 1.**
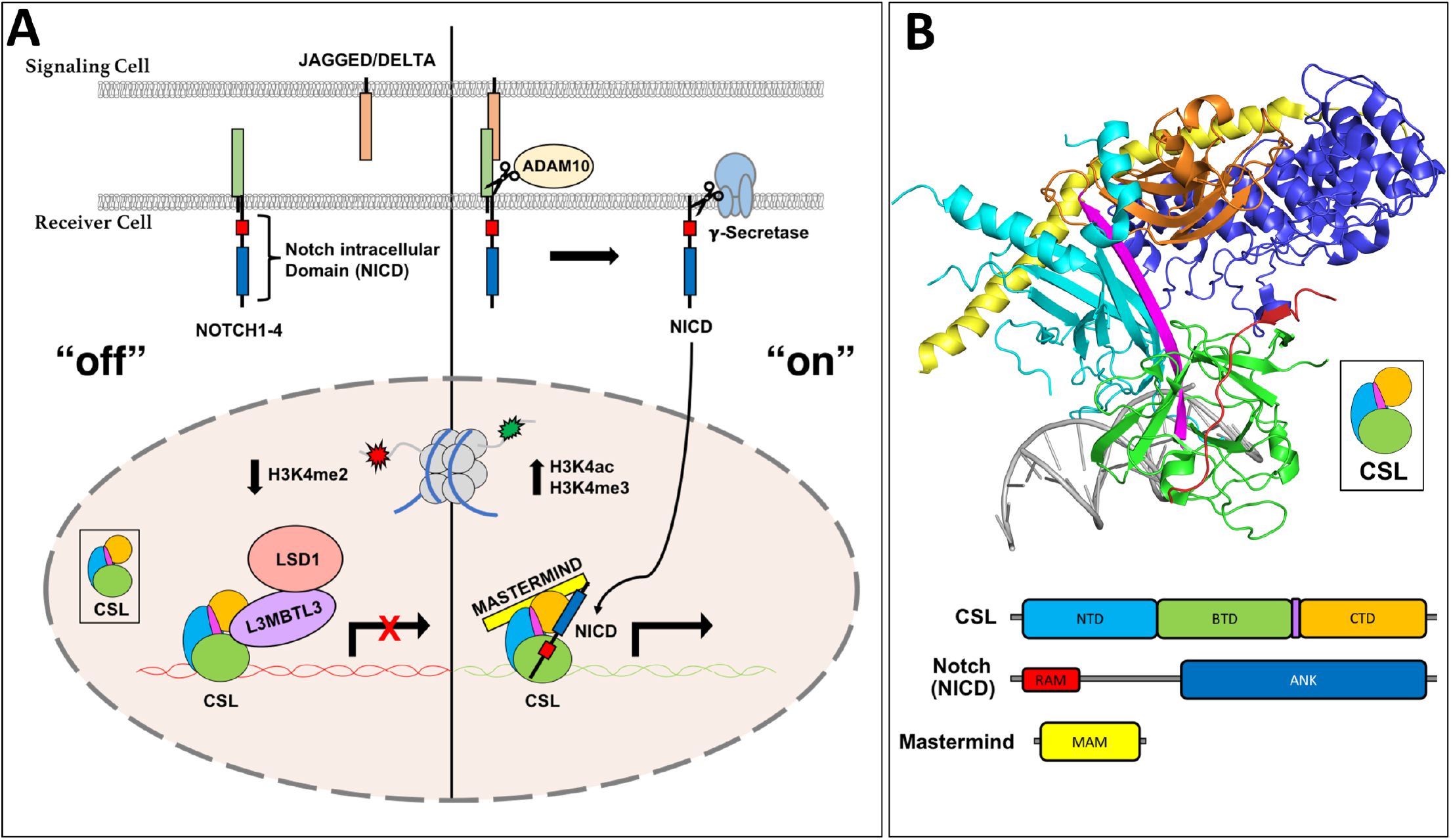
Notch pathway fundamentals. (A) Overview of the Notch signaling pathway. *Left*, “off”: in the absence of Notch receptor-ligand interactions, the transcription factor CSL (RBPJ in mammals) binds corepressor proteins, such as L3MBTL3, which recruits repression machinery to Notch target genes. CSL-L3MBTL3 complexes bind the demethylase LSD1 (KDM1A) leading to a decrease in H3K4me2 epigenetic marks. *Right*, “on”: when the NOTCH receptor is activated by ligand binding, a series of proteolytic events leads to the release of the Notch intracellular domain (NICD), which localizes to the nucleus and forms a ternary activation complex with CSL and Mastermind (MAM). (B) Crystal structure of the CSL-NICD-MAM ternary activation complex bound to DNA (PDBID: 2FO1). The structural core of CSL contains the N-terminal domain (NTD) in cyan, β-trefoil domain (BTD) in green, and C-terminal domain (CTD) in orange, which are integrated into one overall fold by a long β-strand, shown in magenta, that makes hydrogen bonding interactions with all three domains. The RAM (RBPJ Associated Molecule, colored red) domain of NICD forms a high affinity interaction with the BTD of CSL, which tethers the ANK (ankyrin repeats, colored blue) domain nearby. A long kinked α-helix from the MAM N-terminus (yellow) can then bind the ANK-CTD interface and the NTD, locking down the ternary complex.

Mastermind recruits the CDK8 kinase module, which phosphorylates NICD within its PEST domain^8^, leading to its recognition by the E3 ubiquitin ligase FBXW7 (F-box and WD40 repeat protein 7) and ubiquitin-mediated proteasomal degradation of NICD^10,11^. CSL can also function as a repressor by binding to a diverse repertoire of transcriptional corepressors, e.g. in mammals, a specific splice variant of FHL1 (Four and half Lim domains protein 1), also known as KYOT2, RITA1 (RBPJ interacting Tubulin associated protein 1) and SHARP (SMRT/HDAC1 Associated Repressor Protein), also known as MINT or SPEN, that are part of higher order multicomponent repression complexes, which apply repressive marks to histone tails (Fig 1A)^7,12^.

CSL is comprised of three structural domains: the NTD (N-terminal domain), BTD (β-trefoil domain), and CTD (C-terminal domain) (Fig 1B)^13^. These are held in a concise three-dimensional fold by a single β-strand spanning all three domains of the protein (colored magenta in Fig 1B). The RAM (RBPJ associated molecule) domain of NICD forms a high affinity (∼20nM) interaction with an exposed hydrophobic surface on the BTD, tethering the ANK (ankyrin repeats) domain to CSL, which binds only weakly to the CTD^14-16^. Mastermind forms an extended α-helix that binds a composite surface created by ANK and CTD, as well as the NTD^16,17^. Several crystal structures of CSL in complex corepressors have also been solved. For example, in mammals FHL1^18^, RITA1^19^, and SHARP^20^ all have RAM-like motifs that bind to the BTD, illustrating a common binding mode for corepressors to interact with RBPJ (mammalian CSL)^7,12^. More recently, a novel RBPJ-binding corepressor, termed L3MBTL3, was identified in a proteomics screen from a glioma cell line^21^.

L3MBTL3 [Lethal (3) malignant brain tumor-like 3] is a member of the malignant brain tumor (MBT) family of transcriptional repressors that contain between one and four MBT domains (Fig 2A)^22^. These domains impart the ability to bind mono- and dimethylated lysine residues on histone tails, with some MBT proteins showing strict specificity while others, including L3MBTL3, binding promiscuously to methylated lysine residues^23^. Although the precise mechanism of repression is unknown, L3MBTL3 is a putative PcG (Polycomb group) protein that likely facilitates chromatin modification and compaction^24^. More recently, it has become clear that MBT proteins can also recognize methylated lysines on non-histone proteins as well^25,26^. In this role, L3MBTL3 has been shown to function as an adaptor for the CRL4^DCAF5^ E3 ubiquitin ligase, targeting the DNA methyltransferase DNMT1 and the stem cell regulator SOX2 for ubiquitin mediated proteasomal degradation^25^. While all of its roles *in vivo* are still being elucidated, it is known that germline deletion of L3MBTL3 in mice leads to an overabundance of immature erythrocytes, causing embryonic lethality by anemia at E18^27^.

**Figure 2.**
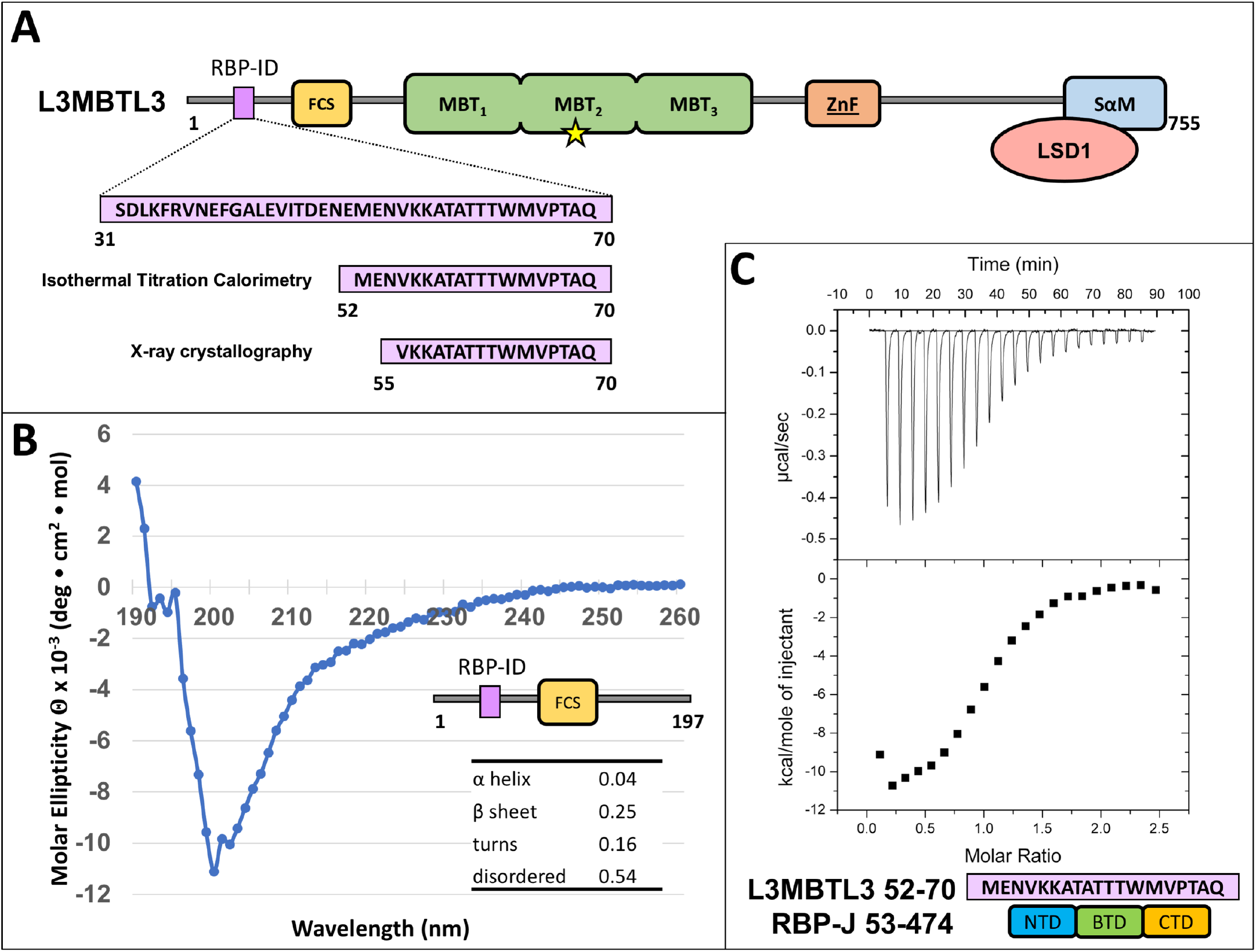
The L3MBTL3 N-terminus contains an RBP-Interaction Domain (RBP-ID). (A) L3MBTL3 domain schematic. L3MBTL3 contains an FCS (phenyalanine-cysteine-serine) zinc finger in yellow, three MBT (malignant brain tumor) methyllysine recognition domains in green, a second canonical zinc finger in orange, and a C-terminal SAM (sterile alpha motif) domain in blue that mediates binding to LSD1/KDM1A (colored salmon). Yellow star denotes Asp 381, which is important for methyllysine binding. The N-terminal RBPJ-Interaction Domain (RBP-ID, colored purple) is ∼15 residues in length with the highlighted 19-mer and 16-mer peptides that were used in the isothermal titration calorimetry (ITC) and X-ray crystallography experiments, respectively. (B) Far UV circular dichroism (CD) of the unbound N-terminal domain of L3MBTL3 (1-197) suggests that this region is mostly random coil with little α-helical structure and modest amounts of β-sheet. (C) Representative ITC binding experiment of RBPJ (53-474) and L3MBTL3 (52-70) peptide shows that it is a 1:1 interaction with 0.92 μM affinity.

As shown in Figure 2A, human L3MBTL3 (isoform b) is a 755 residue multidomain protein comprised of a N-terminal region (∼200 residue) that is predicted to be largely random coil, followed by three MBT domains, a second region of random coil, and a C-terminal SAM (sterile alpha motif) domain. Additionally, there are two predicted zinc finger domains, the FCS-type [phenylalanine (F), cysteine (C), serine (S)] and a classical ZnF type in the N-terminal and C-terminal regions of the protein, respectively. L3MBTL3 binds methylated lysine residues with its second MBT domain and mutation of Asp 381 within this domain (denoted as a yellow star in Fig 2A) has been shown to disrupt interactions with both methyllysine peptides and UNC1215, a potent small molecule inhibitor of L3MBTL3 methyllysine reader function^28,29^. The SAM domain is involved in homo- and heteromultimerization of L3MBTL3^30^.

Previously, we identified an N-terminal region of L3MBTL3 that was required for binding to RBPJ (mouse CSL ortholog) *in vitro* and in cells^21^ (Fig 2A). Here we further define this region and isolate an L3MBTL3 peptide that is necessary and sufficient for interacting with RBPJ. Moreover, we use this information to determine the crystal structure of the RBPJ-L3MBTL3 corepressor complex bound to DNA, and based on the structure, we perform a comprehensive thermodynamic binding analysis. Despite the lack of sequence similarity, we show that L3MBTL3 binds to RBPJ similar to other BTD-binding proteins, such as the RAM domain of NICD or other corepressors, e.g. KYOT2 and RITA1^7,12^; however, L3MBTL3 has an atypical insertion of three threonine residues, resulting in an unusual peptide backbone conformation not seen in other BTD-binding proteins. We validate our structural findings using structure-guided mutants of RBPJ and L3MBTL3, and test these mutants using ITC and cell-based assays. We observe a high degree of correspondence between our structure and the molecular/functional consequences associated with the expression of various RBPJ/L3MBTL3 mutants in mammalian cells. Moreover, to identify potential target genes regulated by RBPJ-L3MBTL3, we perform RNA-Seq on a mouse hybridoma mature T (MT) cell line, in which RBPJ has been depleted using CRISPR-Cas9 technology and replaced by a L3MBTL3-binding-deficient RBPJ mutant. We further validate these findings by shRNA mediated knockdown of L3MBTL3 in MT cells.

## Results

### Defining the RBPJ-Interaction Domain of L3MBTL3

We used isothermal titration calorimetry (ITC) to measure the binding constants between constructs of human L3MBTL3 and mouse RBPJ in order to map the RBPJ-interaction domain (RBP-ID) of L3MBTL3. To begin, we found that an L3MBTL3 construct (Fig 2A and Table 1), which contains its N-terminal region through the MBT domains (residues 1-523), binds RBPJ with micromolar affinity (K_d_ = 1.9 μM). Dividing this construct into two parts – the MBT domains (198-523) were shown to have no detectable binding to RBPJ, while the N-terminus (1-197) bound with a dissociation constant of 1.5μM (Table 1), suggesting that the MBT domains do not contribute to interactions with RBPJ. Circular dichroism (CD) was performed on the N-terminus construct (1-197) to identify any secondary structural elements. In agreement with *in silico* secondary structure predictions (SABLE server for example^31^), the CD data showed that the isolated N-terminal construct is composed of primarily random coil, with some potential β-sheet and very low α-helix content (Fig 2B). Further dissection of the N-terminus led to characterization of L3MBTL3 (31-70) with a K_d_ of 450nM, which we could additionally narrow down to a 19-mer peptide (52-70) that has a comparable affinity (0.92 μM K_d_) (Fig 2C and Table 1). Based on the crystal lattice contacts of previous RBPJ-coregulator X-ray structures^18,19^, we designed a 16-mer peptide (55-70) that was used for crystallization trials and bound RBPJ with a 530 nM K_d_ (Table 1).

**Table 1.** Calorimetric binding data for L3MBTL3 constructs and native RBPJ.

| **L3MBTL3** | **$K$ ($M^{-1}$)** | **$K_d$**<br>**($\mu M$)** | **$\Delta G^\circ$**<br>**($kcal/mol$)** | **$\Delta H^\circ$**<br>**($kcal/mol$)** | **$-T\Delta S^\circ$**<br>**($kcal/mol$)** |
| --- | --- | --- | --- | --- | --- |
| 1-523 | $5.4 \pm 0.9 \times 10^5$ | 1.9 | $-7.7 \pm 0.2$ | $-10.6 \pm 1.6$ | $2.9 \pm 1.8$ |
| 198-523 | NBD | --- | --- | --- | --- |
| 1-197 | $6.6 \pm 0.3 \times 10^5$ | 1.5 | $-8.0 \pm 0.03$ | $-7.2 \pm 0.1$ | $-0.7 \pm 0.1$ |
| 31-70* | $2.3 \pm 0.3 \times 10^6$ | 0.45 | $-8.7 \pm 0.1$ | $-7.5 \pm 0.8$ | $-1.1 \pm 0.8$ |
| 52-70 | $1.1 \pm 0.1 \times 10^6$ | 0.92 | $-8.2 \pm 0.1$ | $-13.1 \pm 0.3$ | $4.8 \pm 0.3$ |
| 55-70 | $2.3 \pm 0.7 \times 10^6$ | 0.53 | $-8.7 \pm 0.2$ | $-8.9 \pm 1.2$ | $0.2 \pm 1.4$ |

All experiments were performed at 25°C. NBD represents no binding detected. Values are the mean of at least three independent experiments and errors represent the standard deviation of multiple experiments. \*Binding data from Xu et al.<sup>21</sup>

### Crystal Structure of the RBPJ-L3MBTL3-DNA Complex

In order to generate crystals of the complex between the structural core of RBPJ (residues 53-474) and the RBP-ID of L3MBTL3, we tested different constructs of RBPJ and different L3MBTL3 peptides, as well as different oligomeric DNA duplexes that contain a single RBPJ binding site. From our previous work demonstrating that L3MBTL3 competes with NICD for RBPJ binding^21^, we surmised that L3MBTL3 in complex with RBPJ may crystallize in conditions similar to other coregulators that have RAM-like peptide binding motifs, *e*.*g*. FHL1 (PDB: 4J2X)^19^ or RITA1 (PDB: 5EG6)^18^. However, RBPJ-L3MBTL3-DNA complexes did not crystallize under these previously identified conditions. Using a different N-terminal affinity tag (His-SMT3 rather than GST), which results in an N-terminus of RBPJ shortened by four residues following cleavage, was critical for crystallizing RBPJ-L3MBTL3-DNA complexes. Additionally, in contrast to previous RBPJ complex structures, which used an oligomeric DNA duplex that corresponds to a RBPJ binding site within the *Hes-1* promoter region^13,14,16,18-20,32^, we used a C→T variant of this sequence (**<u>C</u>**GTGGGAA vs **<u>T</u>**GTGGGAA) that interacts with higher affinity to RBPJ (*data not shown*). Combining these two approaches successfully led to the identification of multiple crystallization conditions for the RBPJ-L3MBTL3 complex bound to DNA and optimization of conditions led to large, X-ray diffraction quality crystals. Molecular replacement performed with the RBPJ-DNA complex (PDB: 3IAG)^33^ was used to solve the initial structure, allowing L3MBTL3 to be built into the F_o_-F_c_ map. Interestingly, the N-terminal serine residue of RBPJ forms a key crystal contact with a neighboring DNA molecule, underscoring the importance of the new RBPJ construct used for structural studies. We report the 2.06 Å resolution RBPJ-L3MBTL3-DNA structure (Fig 3A) from P2_1_2_1_2_1_ crystals (a = 67.9, b = 96.9, c = 105.8) with a single copy of each component in the asymmetric unit. The final dataset was refined to R_work_ and R_free_ values of 19.9% and 24.3%, respectively (Table 2).

**Figure 3.**
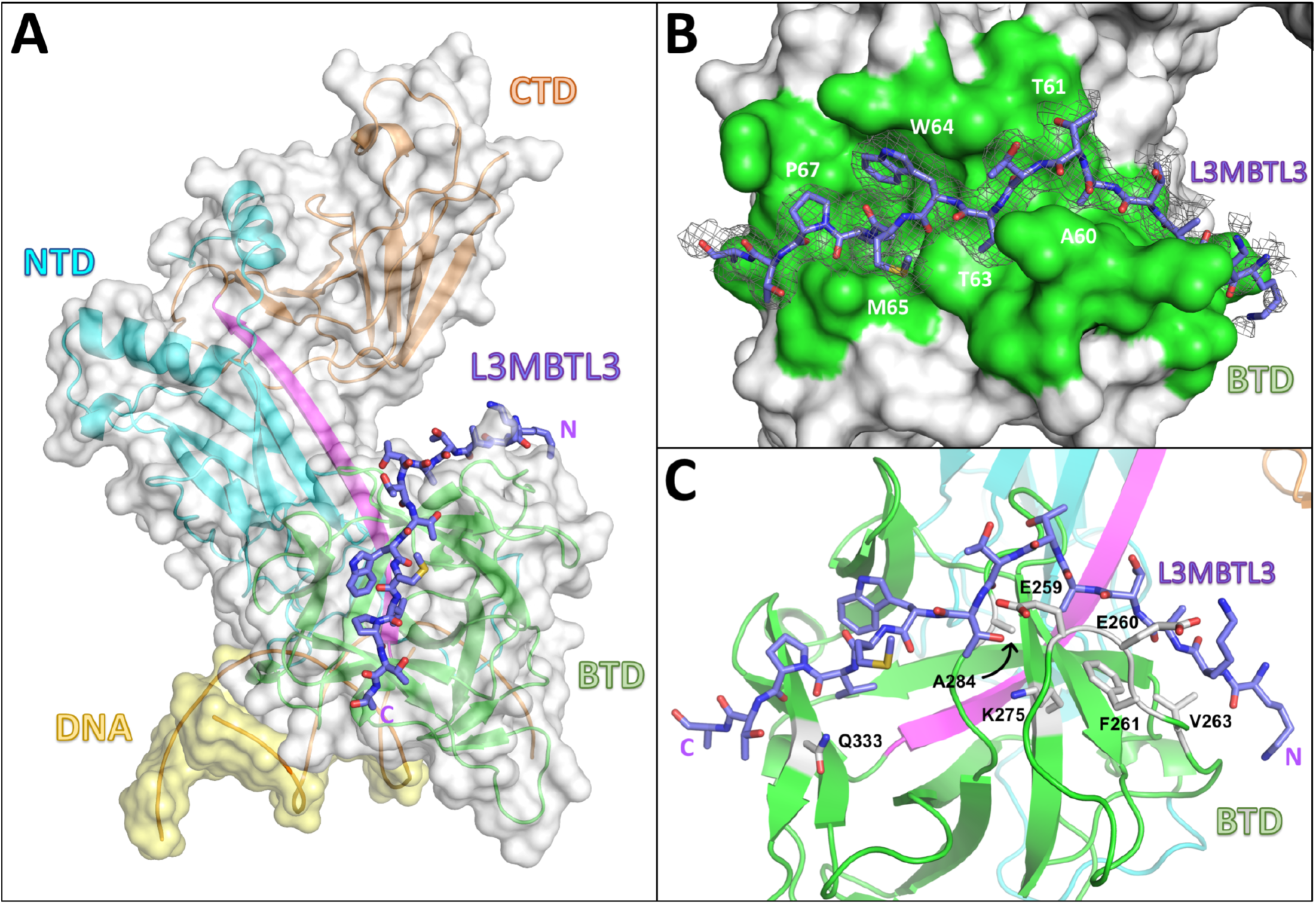
RBPJ-L3MBTL3-DNA Crystal Structure. (A) The RBPJ-L3MBTL3-DNA X-ray structure with the NTD, BTD, and CTD colored cyan, green, and orange, respectively. The DNA wire model is shown in yellow. L3MBTL3 RBP-ID (55-70) represented as purple sticks binds as an elongated peptide along the top and front faces of the BTD. (B) Figure shows the L3MBTL3 binding pocket (colored green) on the BTD of RBPJ. RBPJ residues that directly contact L3MBTL3 were determined by the PISA server^34^. The 2F_o_-F_c_ electron density map contoured at 1σ corresponds to the L3MBTL3 peptide. (C) L3MBTL3 binds residues in the BTD, depicted as grey sticks, that are important for binding RAM and other RAM-like coregulators.

**Table 2.**
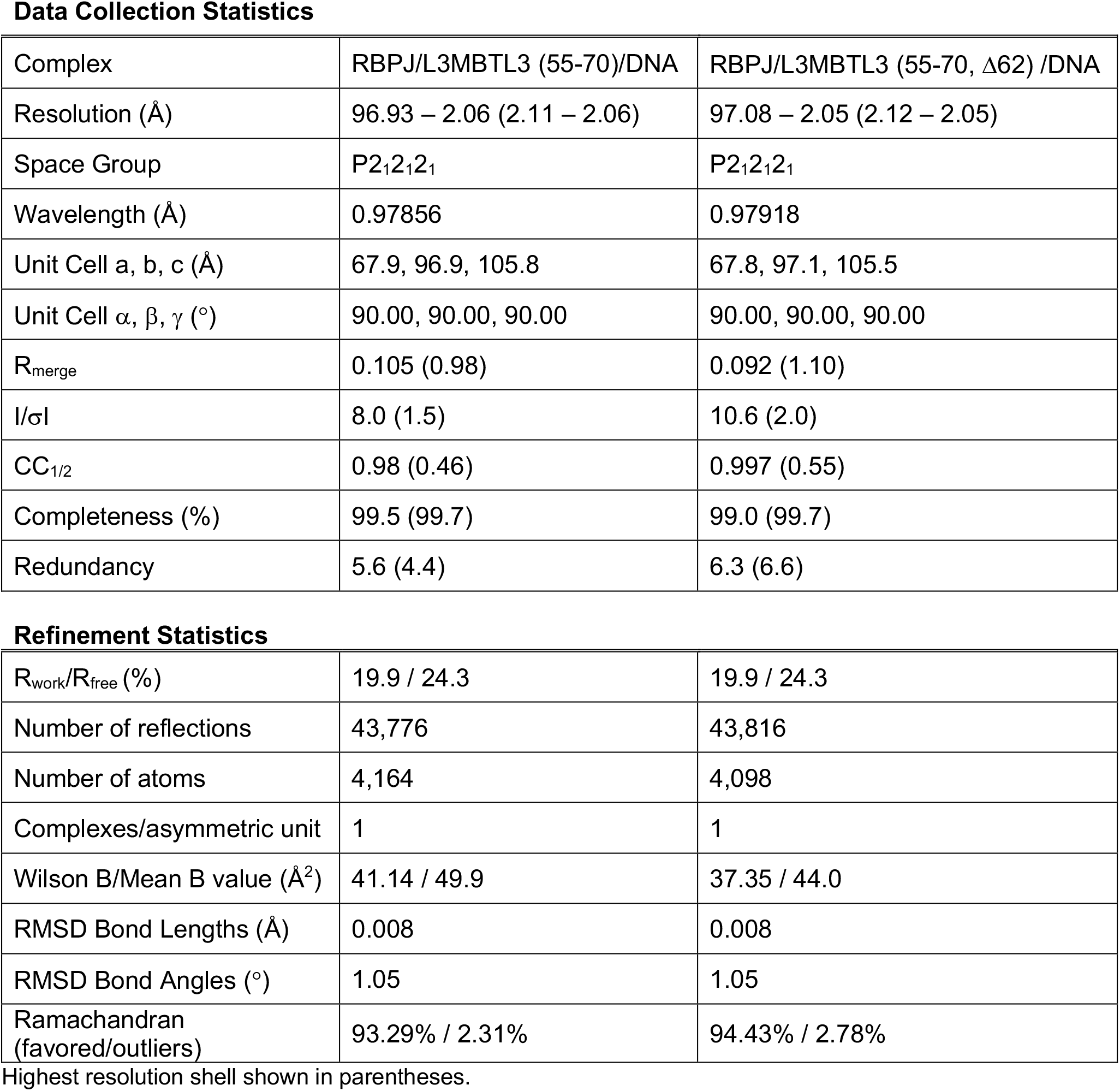
X-ray Data collection and refinement statistics.

|  |  |  |
| --- | --- | --- |
| Complex | RBPJ/L3MBTL3 (55-70)/DNA | RBPJ/L3MBTL3 (55-70, $\Delta 62$) /DNA |
| Resolution (Å) | 96.93 – 2.06 (2.11 – 2.06) | 97.08 – 2.05 (2.12 – 2.05) |
| Space Group | P2<sub>1</sub>2<sub>1</sub>2<sub>1</sub> | P2<sub>1</sub>2<sub>1</sub>2<sub>1</sub> |
| Wavelength (Å) | 0.97856 | 0.97918 |
| Unit Cell a, b, c (Å) | 67.9, 96.9, 105.8 | 67.8, 97.1, 105.5 |
| Unit Cell $\alpha$, $\beta$, $\gamma$ (°) | 90.00, 90.00, 90.00 | 90.00, 90.00, 90.00 |
| R<sub>merge</sub> | 0.105 (0.98) | 0.092 (1.10) |
| I/σI | 8.0 (1.5) | 10.6 (2.0) |
| CC<sub>1/2</sub> | 0.98 (0.46) | 0.997 (0.55) |
| Completeness (%) | 99.5 (99.7) | 99.0 (99.7) |
| Redundancy | 5.6 (4.4) | 6.3 (6.6) |
| **Refinement Statistics** |  |  |
| R<sub>work</sub>/R<sub>free</sub> (%) | 19.9 / 24.3 | 19.9 / 24.3 |
| Number of reflections | 43,776 | 43,816 |
| Number of atoms | 4,164 | 4,098 |
| Complexes/asymmetric unit | 1 | 1 |
| Wilson B/Mean B value (Å<sup>2</sup>) | 41.14 / 49.9 | 37.35 / 44.0 |
| RMSD Bond Lengths (Å) | 0.008 | 0.008 |
| RMSD Bond Angles (°) | 1.05 | 1.05 |
| Ramachandran (favored/outliers) | 93.29% / 2.31% | 94.43% / 2.78% |

Highest resolution shell shown in parentheses.

The structure shows that consistent with our current and previous binding studies^21^ L3MBTL3 binds entirely to the BTD of RBPJ, threading through a narrow groove and down the front face of the BTD (Fig 3A,B). L3MBTL3 residues 56-69 (-KKATATTTWMVPTA-) were built into the electron density; however, the N-terminal lysines (56-57), which contribute to binding (*see below*), have poorly resolved sidechain density (Fig 3B). L3MBTL3 residues W64 and V66 bury their sidechains into an exposed hydrophobic pocket on the surface of the BTD. The overall binding mode of L3MBTL3 is similar to other RAM-like coregulators that bind RBPJ^7,12^, and analysis of the side chains involved in complex formation (PDBePISA server^34^, shown in Fig 3B) suggests the involvement of many RBPJ residues that have been experimentally shown to impact binding by RAM and other RAM-like partners (Fig 3C)^35^.

### Comparison of RBPJ-L3MBTL3 Complex to Other Coregulators

Structural alignment of L3MBTL3 with the RAM domain of Lin-12 and the corepressors FHL1, RITA1, and SHARP, reveals similar and unique features of L3MBTL3 binding (Fig 4). On one hand, the C-terminal portion of the L3MBTL3 peptide binds RBPJ nearly identically to the other coregulators (Fig 4A), including a perfect alignment of W64 (black box in Fig 4B) with the tryptophan that is conserved in RAM and other BTD-binders with the exception of SHARP, which has a serine residue at this position (Fig 4B). This buried tryptophan is part of the hydrophobic tetrapeptide motif (-ΦWΦP-, Φ = nonpolar residue) that has been found in many BTD-binding proteins and is critical for complex formation^7,12^. However, the L3MBTL3 tetrapeptide sequence is -TWMV- making it the most divergent RBPJ binding partner apart from SHARP, which in addition to the BTD also binds the CTD^20^. P67 (blue box Fig 4B), which resides directly after the tetrapeptide, is conserved in the RAM domains from *C. elegans* and NOTCH4, as well as RITA1 and SHARP. In L3MBTL3, this residue plays an important role in complex formation, as mutation of P67 to alanine completely abolishes L3MBTL3 binding to RBPJ (*see below*).

**Figure 4.**
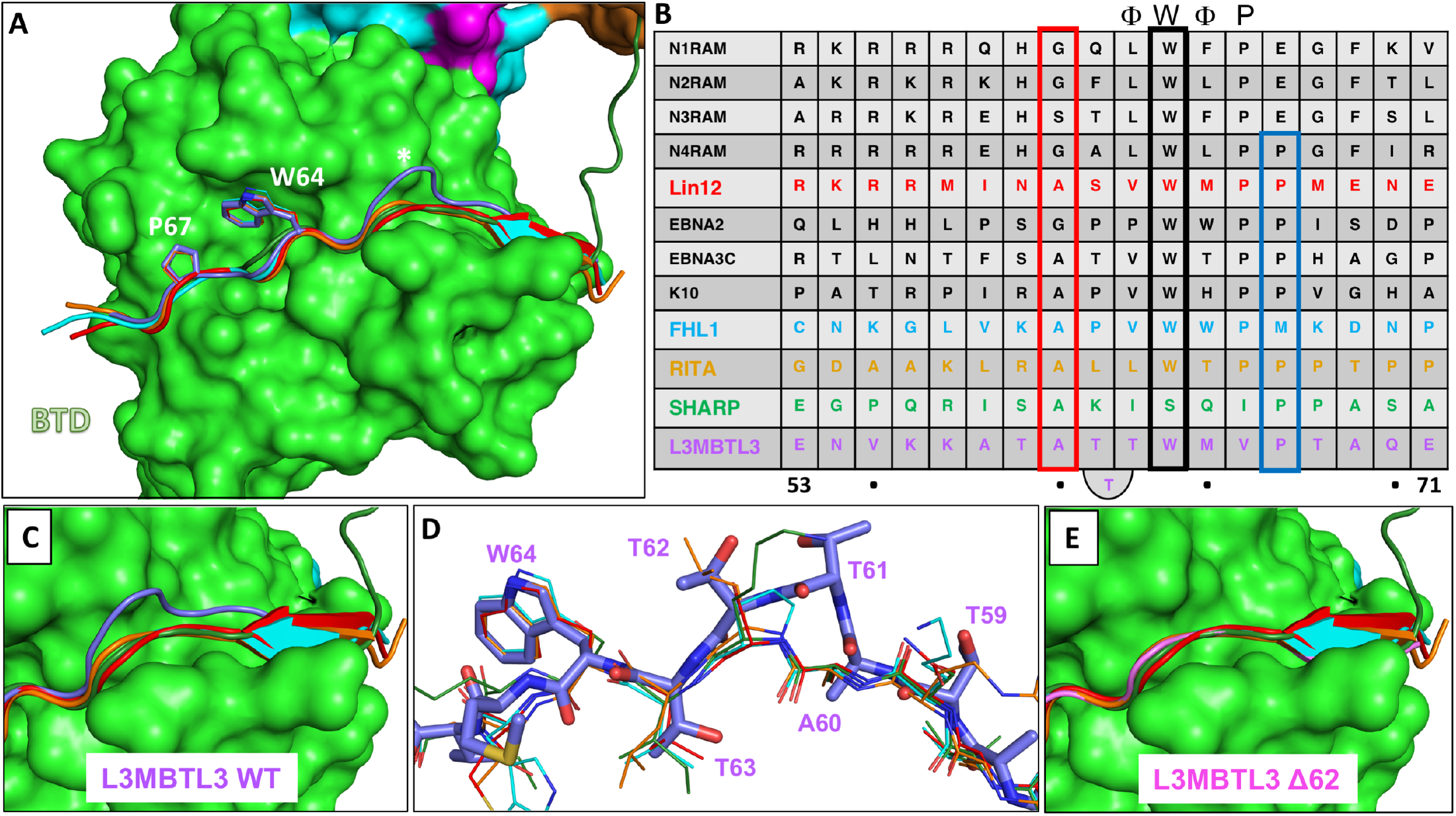
L3MBTL3 Adopts a Distinct Structure Among RBPJ Coregulators. (A) Structural alignment of RAM and RAM-like peptides bound to RBPJ shows a recurrent interaction conformation. RAM from the *C. elegans* Notch ortholog Lin12 (red), FHL1 (light blue), RITA1 (orange), SHARP (dark green), and L3MBTL3 (purple) all bind to the CSL BTD (green surface) as extended peptides. Shown as sticks are L3MBTL3 residues W64 and P67, as well as the corresponding conserved residues in other coregulators. Unlike other coregulators, the backbone of L3MBTL3 (purple) diverges structurally by forming a large bulge over the cavity joining the top and front of the BTD, which is denoted with an asterisk. (B) Structure based sequence alignment of L3MTBL3 with other BTD binding proteins. The conserved tryptophan residue of the hydrophobic tetrapeptide, which has the sequence -ΦWΦP- where Φ = hydrophobic residue, is boxed in black. In contrast to other coregulators, L3MBTL3 has a valine in the fourth position of the -ΦWΦP- instead of a proline, but has a proline immediately following this position (blue rectangle), which is conserved in RITA1, SHARP, and some Notch receptor orthologs. A60 aligns to the strongly conserved position, which requires small sidechain residues (red rectangle); however, this leads to L3MBTL3 having three threonine residues between the alanine and tryptophan, whereas all other coregulators only have two residues. (C) Top-down view of the RBP-ID N-terminus showing that the L3MBTL3 extrusion (purple) pushes its backbone out of alignment with respect to the other coregulators. (D) Aligned stick models of RAM-like peptides show the conserved positions of L3MBTL3 A60 and W64. T63 essentially realigns with the first hydrophobic tetrapeptide residues, forcing the T61/T62 dithreonine to create the structural bulge. (E) The crystal structure of L3MBTL3 Δ62 peptide (pink) bound to RBPJ demonstrates a complete realignment of L3MBTL3 with the other RAM-like peptides, in which L3MBTL3 adopts the β-strand interaction seen in RAM, FHL1, and RITA1.

On the other hand, L3MBTL3 residues upstream of the hydrophobic tetrapeptide differ significantly from the other BTD-binders (Fig 4C). RAM, FHL1, and RITA1 form β-sheet interactions with the BTD and then lie along a groove that runs across the front face of the BTD. The L3MBTL3 N-terminus is situated further inward before its backbone bulges out and over the BTD groove (Fig 4C-D). The extrusion is formed by three consecutive threonines (T61-T63), whereby T63 comes back into alignment with other BTD-binders at the putative start of the L3MBTL3 hydrophobic tetrapeptide (-TWMV-). Before the threonine extrusion, L3MBTL3 A60 aligns with another conserved position among BTD binders (red box in Fig 4B). This position requires residues with a small sidechain due to its close contact with the BTD (Fig 4D). Thus, A60 and T63 anchor L3MBTL3 to the BTD in two places and necessitate an extrusion of a short loop by the intervening two threonine residues. As depicted in Figure 4B, compared to other BTD binders, L3MBTL3 essentially contains an extra residue that imparts this feature in the complex structure.

To further characterize the unusual threonine loop, we reproduced our crystals with an L3MBTL3 peptide in which T62 was deleted (Δ62: VKKATATTWMVPTAQ). The data quality from the L3MBTL3 mutant crystals was nearly identical to the wild-type crystals and the structures refined similarly (Table 2). With one less residue between A60 and T63, the Δ62 peptide takes on a conformation very similar to the other BTD-binders, rather than maintaining the extruded loop (Fig 4E). Notably, the N-terminus of the peptide forms a β-strand interaction akin to RAM, FHL1, and RITA. Moreover, the N-terminal lysine sidechains are also resolved in this structure: K56 extends out into the solvent and K57 rests on the BTD with its amino group making a likely hydrogen bonding interaction with the hydroxyl group from T262 of RBPJ, consistent with this residue providing an important contact between the proteins. Binding of the Δ62 L3MBTL3 peptide to RBPJ (0.99 μM K_d_) is similar to wild-type (0.92 μM K_d_) (Table 3). While the functional significance of the threonine bulge requires further study, retaining an extra threonine raises the possibility of post-translational modification of L3MBTL3 for regulated binding to RBPJ, albeit to date there is no experimental evidence that any of the threonine residues are modified *in vivo*.

**Table 3.** Calorimetric binding data for L3MBTL3 alanine mutants and native RBPJ.

| L3MBTL3 | $K$ ($M^{-1}$) | $K_d$<br>($\mu M$) | $\Delta G^\circ$<br>(kcal/mol) | $\Delta H^\circ$<br>(kcal/mol) | $-T\Delta S^\circ$<br>(kcal/mol) | $\Delta\Delta G^\circ$<br>(kcal/mol) |
| --- | --- | --- | --- | --- | --- | --- |
| WT | $1.1 \pm 0.1 \times 10^6$ | 0.92 | $-8.2 \pm 0.1$ | $-13.1 \pm 0.3$ | $4.8 \pm 0.3$ | -- |
| WT + DMSO | $1.1 \pm 0.1 \times 10^6$ | 0.90 | $-8.2 \pm 0.1$ | $-12.8 \pm 0.6$ | $4.5 \pm 0.7$ | -0.02 |
| N54A | $2.1 \pm 0.6 \times 10^6$ | 0.47 | $-8.6 \pm 0.2$ | $-11.1 \pm 0.9$ | $2.5 \pm 1.1$ | -0.4 |
| V55A | $1.1 \pm 0.2 \times 10^6$ | 0.87 | $-8.3 \pm 0.1$ | $-13.6 \pm 1.0$ | $5.3 \pm 1.1$ | -0.02 |
| K56A + DMSO | $5.6 \pm 1.8 \times 10^5$ | 1.8 | $-7.8 \pm 0.2$ | $-8.0 \pm 1.2$ | $0.2 \pm 1.3$ | 0.4* |
| K57A | $1.9 \pm 0.4 \times 10^5$ | 5.3 | $-7.2 \pm 0.2$ | $-11.9 \pm 0.7$ | $4.7 \pm 0.7$ | 1.1 |
| A58R | $1.9 \pm 0.3 \times 10^6$ | 0.54 | $-8.6 \pm 0.1$ | $-14.0 \pm 0.9$ | $5.4 \pm 0.9$ | -0.3 |
| T59A | $7.6 \pm 1.1 \times 10^5$ | 1.3 | $-8.0 \pm 0.1$ | $-9.9 \pm 0.7$ | $1.9 \pm 0.8$ | 0.2 |
| A60R | $2.4 \pm 0.5 \times 10^5$ | 4.2 | $-7.3 \pm 0.1$ | $-15.7 \pm 0.3$ | $8.4 \pm 0.4$ | 0.9 |
| T61A | $5.8 \pm 0.6 \times 10^5$ | 1.7 | $-7.9 \pm 0.1$ | $-11.4 \pm 0.6$ | $3.5 \pm 0.6$ | 0.4 |
| T62A | $3.3 \pm 0.5 \times 10^6$ | 0.30 | $-8.9 \pm 0.1$ | $-13.7 \pm 0.3$ | $4.8 \pm 0.4$ | -0.7 |
| $\Delta 62$ | $1.0 \pm 0.03 \times 10^6$ | 0.99 | $-8.2 \pm 0.0$ | $-12.7 \pm 0.7$ | $4.5 \pm 0.7$ | 0.04 |
| T63A | NBD | --- | --- | --- | --- | --- |
| W64A | NBD | --- | --- | --- | --- | --- |
| M65A | $1.0 \pm 0.3 \times 10^6$ | 1.0 | $-8.2 \pm 0.2$ | $-12.4 \pm 1.7$ | $4.3 \pm 1.9$ | 0.1 |
| V66A | NBD | --- | --- | --- | --- | --- |
| P67A | NBD | --- | --- | --- | --- | --- |
| T68A | $8.7 \pm 1.0 \times 10^5$ | 1.2 | $-8.1 \pm 0.1$ | $-11.7 \pm 0.7$ | $3.6 \pm 0.7$ | 0.1 |
| A69R | $6.1 \pm 0.7 \times 10^5$ | 1.63 | $-7.9 \pm 0.1$ | $-13.8 \pm 1.1$ | $5.9 \pm 1.2$ | 0.3 |
| Q70A | $1.1 \pm 0.4 \times 10^6$ | 0.94 | $-8.2 \pm 0.3$ | $-14.2 \pm 0.3$ | $5.9 \pm 0.3$ | 0.01 |

Constructs: L3MBTL3 (52-70) and RBPJ (53-474). All experiments were performed at 25°C. NBD represents no binding detected. Values are the mean of at least three independent experiments and errors represent the standard deviation of multiple experiments. \*Relative to WT + DMSO value. K56A peptide required dissolution in a phosphate buffer with 5% DMSO.

### Binding Analysis of RBPJ-L3MBTL3 Mutants

Next, we used ITC to further understand the molecular determinants of L3MBTL3 binding to RBPJ. To this end we designed a series of point mutants scanning along the L3MBTL3 52-70 peptide, starting with N54, whereby each residue was changed to an alanine, except for native alanines in L3MBTL3, which were changed to arginines (Fig 5A). Overall, the results show high variability in binding with single mutations across the L3MBTL3 RBP-ID (Fig 5B and Table 3). Of the 17 mutants, nine (K56A, K57A, A60R, T61A, T63A, W64A, V66A, P67A, and A69R) reduce binding by approximately 50% or more, with T63A, W64A, V66A, and P67A completely abrogating binding. These latter residues (-**TW**M**VP**-) constitute the hydrophobic tetrapeptide region (-ΦWΦP-) with P67 directly following the hydrophobic tetrapeptide, which is conserved in RITA1, SHARP, and other coregulators (Fig 4B). T63, W64, and V66 sample the hydrophobic binding pocket on the face of the BTD that is used by other corepressors, highlighting a recurring Notch coregulator binding mechanism to RBPJ. The pyrrolidine ring of P67 points away from the BTD face, but its orientation allows for hydrogen bonding and hydrophobic interactions with the backbone of RBPJ. The N-terminal lysines (K56/57) putatively interact with a negative patch on the BTD, and as mentioned above, K57 appears to have specific contacts with T262 of RBPJ while K56 points outwards to the solvent. Under more physiological ionic strength conditions, K56 is likely to sit on the BTD itself and have specific interactions as well. A60R also reduces binding by 75%, primarily due to the steric clashes caused by introduction of the large arginine sidechain where normally the methyl group of alanine points directly towards the surface of the BTD. Unexpectedly, three L3MBTL3 point mutants increase binding to RBPJ by varying degrees. A58R introduces a charged residue near E260 of RBPJ, which has been demonstrated to be a mediator of salt bridge bond formation for FHL1 and RITA1^18,19^. The most outstanding binding increase (300 nM K_d_) is seen for T62A, which is the second residue of the three threonine extrusion in wild-type L3MBTL3 and perhaps reduces some of the backbone strain induced by the loop bulging.

**Figure 5.**
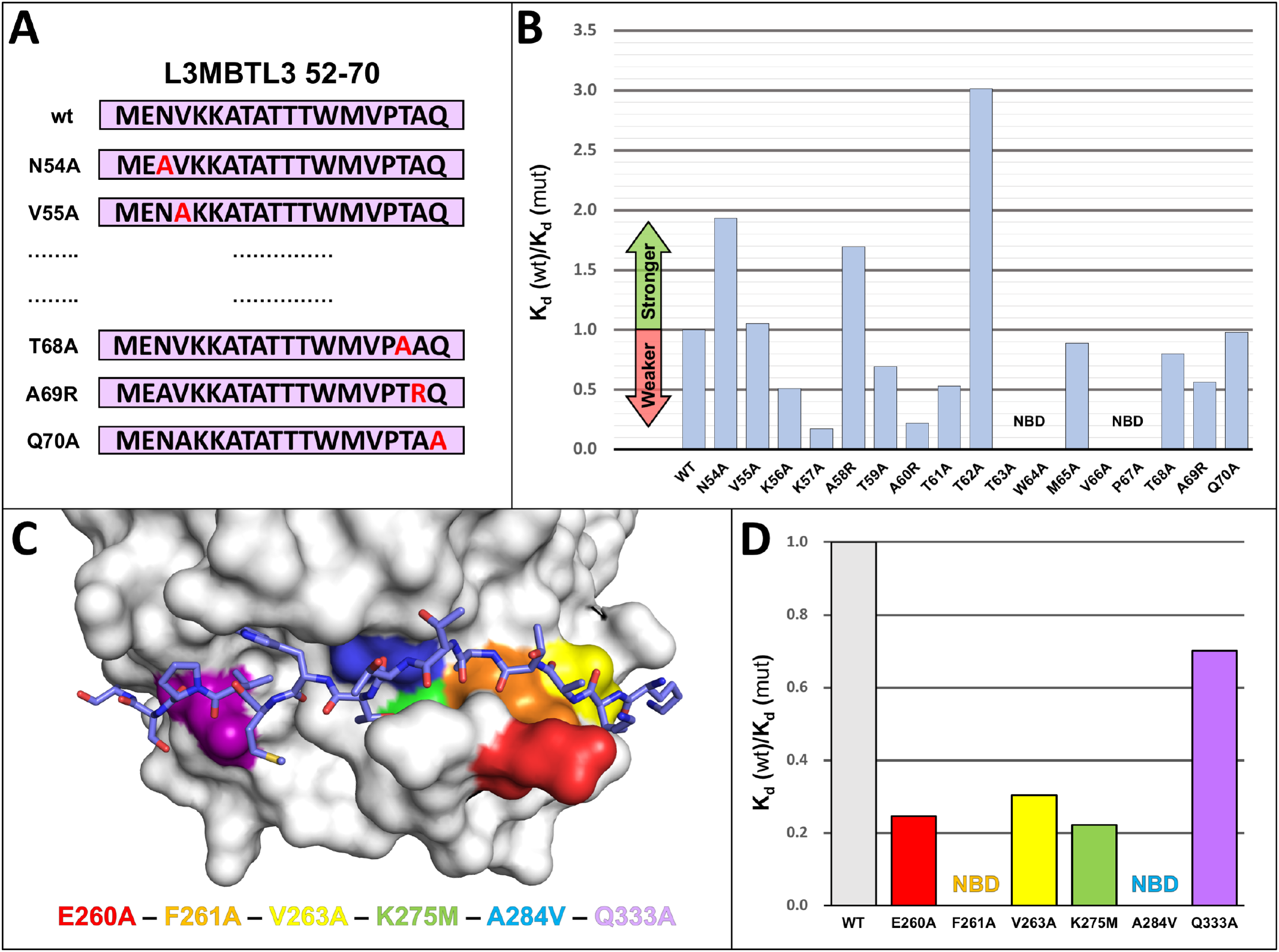
RBPJ-L3MBTL3 Binding Analysis Reveals Residues Sensitive to Mutation. (A) Diagram of scanning point mutants of the 19-mer L3MBTL3 RBP-ID (52-70) for ITC binding studies. Starting at N54, each residue was mutated individually to an alanine, while native alanines were mutated to arginine. (B) Affinity change plot [K_d_ (wt)/K_d_ (mut)] for L3MBTL3 alanine scanning mutants tested against RBPJ wild-type (WT). Changes are plotted as the ratio of K_d_ values, where an increase in K_d_ (weaker binding) is below 1 and vice versa. Mutations along the length of the peptide have varying effects on binding, with K57A and A260R significantly affecting binding; T63A, W64A, V66A, from the hydrophobic tetrapeptide region, and the adjacent P67A mutant all completely abrogate binding. N54A, A58R, and T62A show increased binding to RBPJ, with T62A from the dithreonine loop behaving as the tightest binding peptide (∼300nM K_d_). (C) Visualization of BTD residues targeted for mutation with color coding. The L3MBTL3 peptide is colored purple and shown as sticks. These BTD residues (E260, F261, V263, K275, A284, Q333) have all been shown to influence binding of other RAM-like coregulators, with F261 and A284 mutations causing the largest decreases in affinity. (D) Affinity change plot for BTD mutants tested against L3MBTL3 52-70 wild-type peptide. All mutations negatively affect binding, with Q333A having the smallest effect; F261A and A284V completely abrogate L3MBTL3 binding; and E260A, V263A, and K275M have modest effects on binding. NBD = no binding detected.

We then measured binding of the wild-type L3MBTL3 peptide against a series of RBPJ mutants (Fig 5C and Table 4) known to affect binding of other BTD interacting partners: E260A, F261A, V263A, K275M, A284R, and Q333A^35^. These residues are generally surface exposed that span the path of the coregulator binding site and whose mutation doesn’t alter the overall structure of the BTD. As with the L3MBTL3 point mutants, these mutants again demonstrate the remarkable sensitivity of the RBPJ-L3MBTL3 interaction at certain sites (Fig 5D). In addition to complete disruption of binding by F261A and A284V mutations, E260A (3.73 μM), V263A (3.03 μM), and K275M (4.15 μM) all show a modest ∼3-5 fold increase in K_d_, *i*.*e*. decrease in affinity. Q333A had a small reduction in binding (1.31 μM), which is consistent with this residue having the most variable effect on binding of other coregulators^18-20^. The pattern of L3MBTL3 binding to these mutant forms of RBPJ is consistent with the other BTD-binders, although complete loss of binding by F261 or A284 is unusual^7,12^.

**Table 4.** Calorimetric binding data for native L3MBTL3 and RBPJ mutants.

| | RBPJ | $K$ ($M^{-1}$) | $K_d$<br>($\mu M$) | $\Delta G^\circ$<br>(kcal/mol) | $\Delta H^\circ$<br>(kcal/mol) | $-T\Delta S^\circ$<br>(kcal/mol) | $\Delta\Delta G^\circ$<br>(kcal/mol) |
| --- | --- | --- | --- | --- | --- | --- | --- |
| | WT | $1.1 \pm 0.1 \times 10^6$ | 0.92 | $-8.2 \pm 0.1$ | $-13.1 \pm 0.3$ | $4.8 \pm 0.3$ | -- |
| BTD<br>mutants | E260A | $2.6 \pm 0.8 \times 10^5$ | 3.8 | $-7.4 \pm 0.2$ | $-13.6 \pm 3.4$ | $6.2 \pm 3.6$ | 0.9 |
| | E260K* | $2.2 \pm 0.1 \times 10^5$ | 4.5 | $-7.3 \pm 0.1$ | $-16.8 \pm 0.9$ | $9.5 \pm 0.9$ | 0.9 |
|  | F261A | NBD | --- | --- | --- | --- | --- |
| | V263A | $7.6 \pm 1.5 \times 10^5$ | 1.3 | $-8.0 \pm 0.1$ | $-10.6 \pm 1.0$ | $2.6 \pm 0.9$ | 0.2 |
| | K275M | $2.4 \pm 0.6 \times 10^5$ | 4.2 | $-7.3 \pm 0.1$ | $-13.7 \pm 1.4$ | $6.4 \pm 1.5$ | 0.9 |
|  | A284V | NBD | --- | --- | --- | --- | --- |
| | Q333A | $3.3 \pm 0.9 \times 10^5$ | 3.0 | $-7.5 \pm 0.2$ | $-10.3 \pm 2.1$ | $2.8 \pm 2.3$ | 0.7 |

Constructs: L3MBTL3 (52-70) and RBPJ (53-474). All experiments were performed at 25°C. NBD represents no binding detected. \*N=2 replicates. Values are the mean of at least three independent experiments and errors represent the standard deviation of multiple experiments.

### Cellular Analysis of the RBPJ-L3MBTL3 Complex

To validate in cells the findings from our crystal structure and ITC binding studies, we first tested the activity of L3MBTL3 mutants in coimmunoprecipitation (coIP) and mammalian two-hybrid (MTH) assays. We focused on the residues 63-67 (-TWMVP-), which encompass the hydrophobic tetrapeptide and flanking conserved regions, and had the strongest effect on binding to RBPJ in our ITC experiments. Here, we generated single alanine point mutants in full-length L3MTBL3 that correspond to those tested in an L3MBTL3 peptide ITC, as well as a 5xA mutant with all five residues mutated in tandem (TWMVP → AAAAA). Immunoprecipitation of HA-tagged L3MBTL3 variants from a U87 human glioblastoma cell line (Fig 6A) shows a significant reduction in binding to endogenous RBPJ for all of the mutants except for M65A, whose partial impairment is in accord with the slight increase in K_d_ measured *in vitro*. As a control, the Δ(1-64) L3MBTL3 mutant was previously shown to abrogate binding to RBPJ in cells^21^, which we now recognize as having a truncated RBP-ID. We then moved to an established mammalian two-hybrid (M2H) assay in HeLa cells^36^. In this experiment, L3MBTL3 variants are fused to a Gal4 DNA-binding domain and RBPJ wild-type is fused to the VP-16 activation domain such that RBPJ-L3MBTL3 interactions lead to induction of a luciferase reporter (Fig 6B). As expected, L3MBTL3 wild-type shows a concentration dependent increase in luciferase activity. Consistent with our ITC and coIP assays, the M65A mutant has a negligible decrease in reporter activity; whereas, T63A, W64A, V66A, P67A, and 5xA mutants all severely blunt induction of the luciferase reporter. Taken together, these cellular assays support our structural and binding studies and elucidate the key residues involved in RBPJ-L3MBTL3 complex formation.

**Figure 6.**
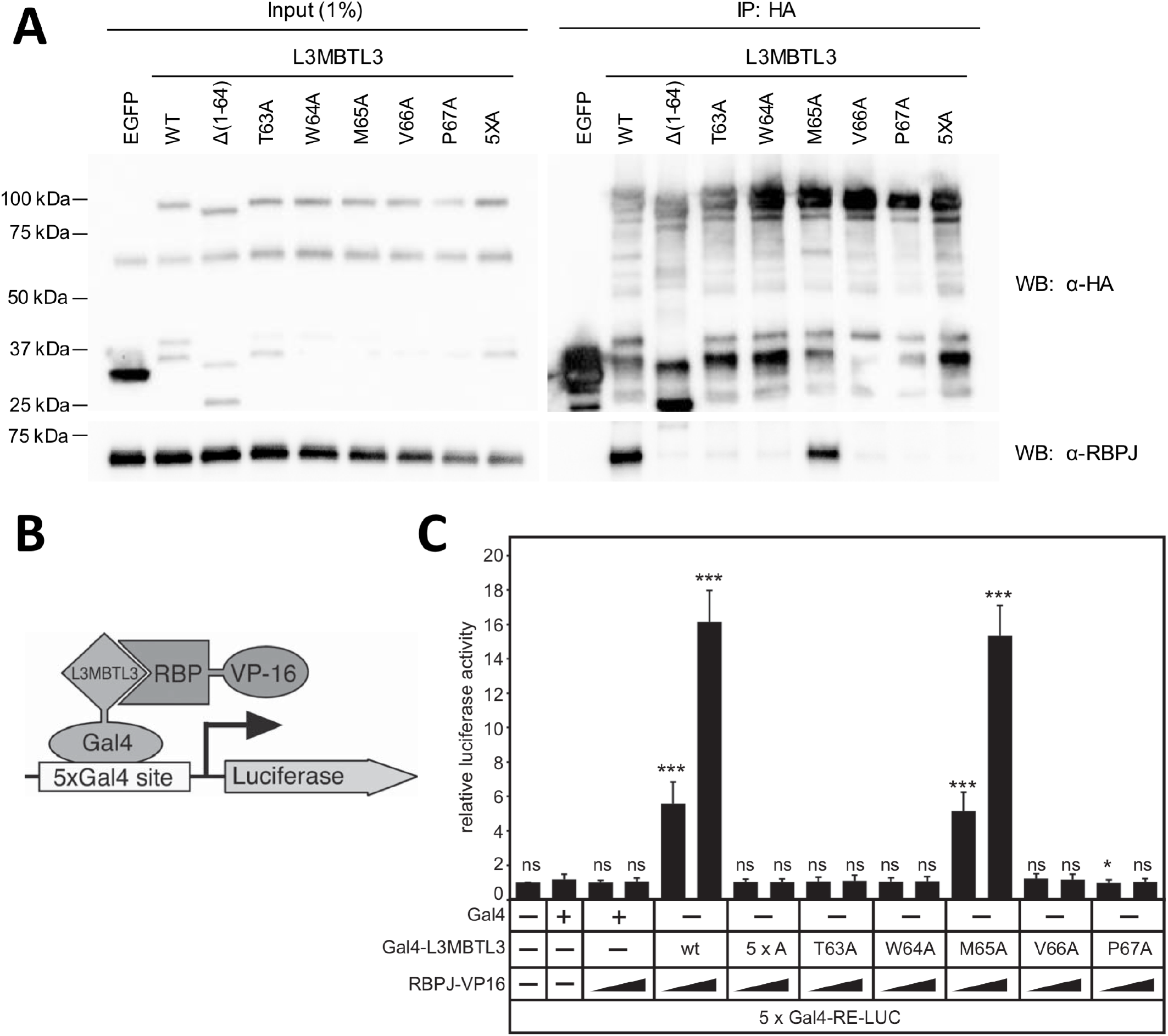
Cellular Analysis of L3MBTL3 Mutants. (A) Figure shows Western blot (WB) of immunoprecipitated (IP) HA-tagged L3MBTL3, wild-type (WT) and mutants, with RBPJ from U87-MG glioblastoma cells. The N-terminal deletion construct L3MBTL3 (Δ1-64) has previously been shown to completely abrogate interactions with RBPJ in cells^21^. The binding deficient L3MBTL3 point mutants identified by ITC (T63A, W64A, V66A, P67A) are similarly impaired for binding RBPJ in cells; whereas, L3MBTL3 M65A, which retains 90% of binding *in* vitro, only modestly affects interactions with RBPJ in cells. As expected, the 5x alanine mutant (5XA = T63A/W64A/M65A/V66A/P67A) completely abrogates L3MBTL3-RBPJ interactions. (B) Schematic representation of mammalian two-hybrid assay. HeLa cells were cotransfected with Gal4-L3MBTL3 wild-type and mutant constructs and increasing amounts of RBP-VP16 together with pFR-Luc containing Gal4 recognition sites (5xGal4-RE-LUC). Luciferase activity was determined from 100 µg portions of total cell extract. Fold-activation was determined by the relative luciferase activity after cotransfection of the Gal4 construct alone. Mean values and standard deviations from four experiments are shown. (C) Gal4-L3MBTL3 and RBPJ-VP16 interact to induce luciferase expression in an RBPJ-VP16 concentration dependent manner. Luciferase expression in the mutants corroborates the binding data and coimmunoprecipitation results, with only the M65A mutant able to recruit RBPJ-V16 to activate the reporter.

To further investigate the biological role of the RBPJ-L3MBTL3 complex in cells, we made use of a mature T (MT) cell line in which Notch is in the OFF state^20^ (Fig 7). In this system, CRISPR/Cas9-mediated depletion of RBPJ leads to upregulation of Notch target genes, due to derepression, and this phenotype can be efficiently rescued by reintroducing wild-type (WT) RBPJ expression^20^ (Fig 7B). Based on the structure and binding studies of the RBPJ-L3MBTL3 complex, we generated RBPJ F261A and A284V single mutants, as well as a F261A/A284V double mutant, and expressed these proteins in MT cells depleted for RBPJ. Western blot (WB) analysis demonstrated that the RBPJ mutants were expressed at similar levels as WT RBPJ (Fig 7A). While WT RBPJ efficiently downregulates the expression of the Notch target genes Lgmn, Hes1 and Hey1 (Fig 7B and as previously described^20^), this was not the case for RBPJ mutants F261A, A284V, and F261A/A284V (Fig 7B), which were defective in repression, supporting our biophysical, biochemical and reporter-based data.

**Figure 7.**
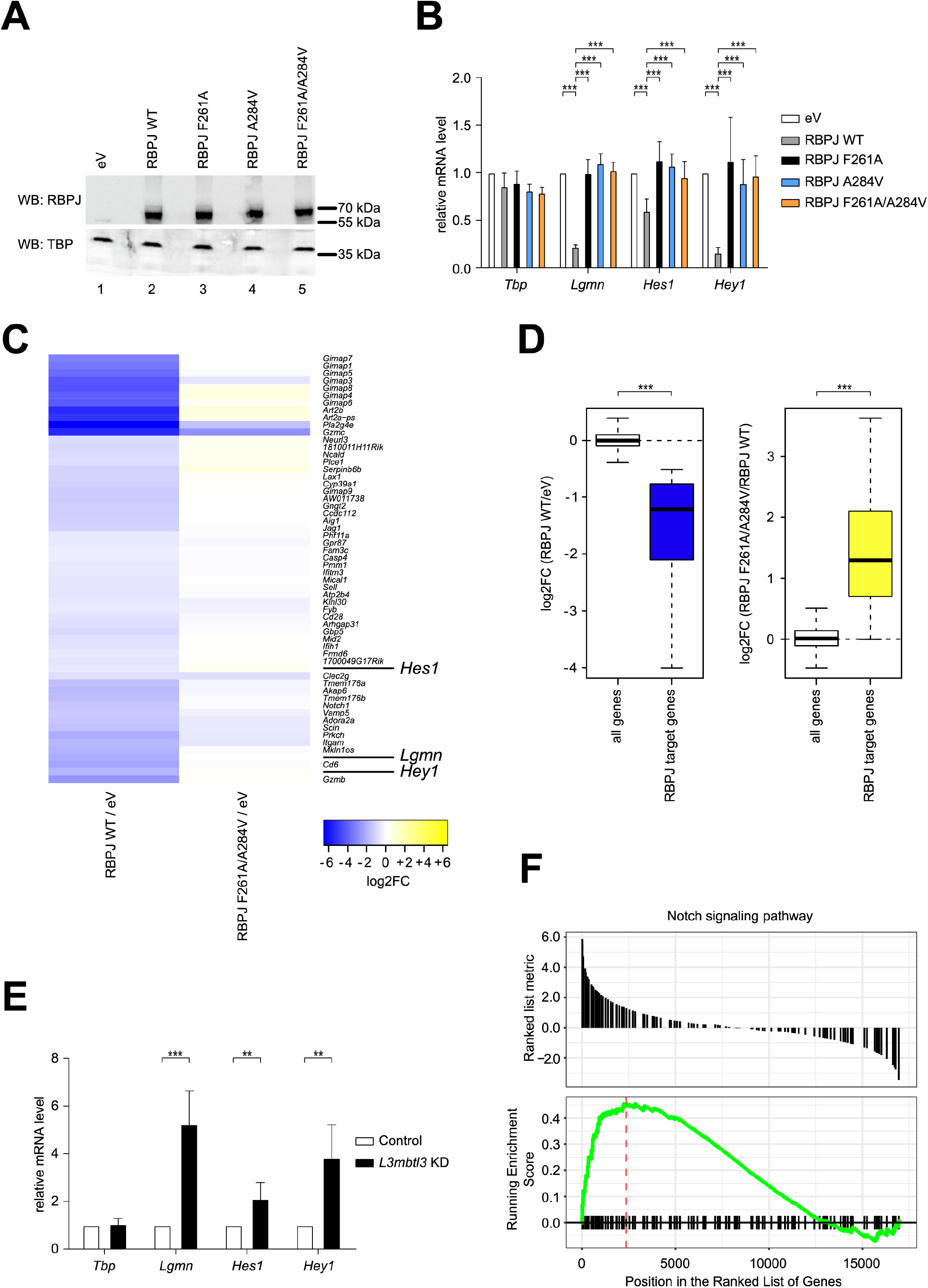
Characterization of RBPJ/L3MBTL3 Interactions in Mature T (MT) Cells. (A) RBPJ wild type (WT) and single or double mutants are efficiently expressed in mature T (MT) cells depleted of endogenous RBPJ. MT cells depleted of RBPJ were infected with viruses carrying plasmids encoding for RBPJ WT, F261A, A284V, F261A/A284V or empty vector (eV) as control. Nuclear extracts were analyzed by Western blotting (WB) with the indicated antibody. TBP was used as loading control. (B) Expression of Notch target genes is downregulated by RBPJ WT but not by the single or double mutants in MT cells depleted of endogenous RBPJ. MT cells depleted of RBPJ were infected with viruses carrying plasmids encoding for RBPJ WT, F261A, A284V, F261A/A284V or empty vector (eV) as control. Upon RNA extraction and reverse transcription, cDNAs were analyzed by qPCR using assays specific for *Tbp, Lgmn, Hes1* or *Hey1*. Data were normalized versus the housekeeping gene *GusB*. The mean ± SD of seven experiments is shown (***p < 0.001, unpaired Student’s t test). (C) Heat map visualization of log2-transformed gene expression changes of genes that are significantly downregulated (FDR < 0.05 and log2FoldChange < - 0.5) upon rescue with RBPJ WT (RBPJ WT / eV) in MT cells depleted of endogenous RBPJ (left panel). Right panel shows the expression changes of these genes upon rescue with the F261A/A284V double mutant (RBPJ F261A/A284V / eV). MT cells depleted of RBPJ were infected with viruses carrying plasmids encoding for RBPJ WT, F261A/A284V or empty vector (eV) as control. Upon RNA extraction samples were analyzed by RNA-Seq. (D) Left: Box plot representation of the log2-transformed gene expression changes (RBPJ WT / eV) from all genes and RBPJ target genes defined as those genes downregulated upon rescue with RBPJ WT. Right: log2-transformed gene expression changes of all genes (RBPJ F261A/A284V / RBPJ WT) and the RBPJ target genes (***p < 0.001, Wilcoxon rank sum test). (E) L3MBTL3 depletion in MT cells leads to upregulation of Notch target genes. MT cells were infected with hairpin directed against L3MBTL3 or scramble (Control) as control. Upon RNA extraction and reverse transcription, cDNAs were analyzed by qPCR using assays specific for *Tbp, Lgmn, Hes1* or *Hey1*. Data were normalized versus the housekeeping gene *GusB*. Shown is the mean ± SD of five experiments (**p < 0.01, ***p < 0.001, unpaired Student’s t test). (F) Visualization of the gene set enrichment analysis (GSEA) comparing L3MBTL3 KD and the scramble (Control) sample indicating significant and concordant induction of genes belonging to the GO term (biological process) “Notch signaling pathway” (adjusted p-value = 0.0128).

To further characterize the effects of the RBPJ-L3MBTL3 interaction on a transcriptomic level, we performed RNA-Seq analysis focusing on MT cells that express either WT RBPJ or the RBPJ double mutant F261A/A284V (Figs 7C-D, S1A-B and Tables S1-S3). The individual replicates showed good reproducibility (Fig S1A) and we were able to detect a group of genes that were significantly downregulated upon rescue with WT RBPJ (Fig 7C-D). Interestingly, the majority of these genes were not downregulated using the RBPJ F261A/A284V mutant (Fig 7C-D). To note, gene ontology (GO) analysis based on the significantly downregulated genes upon RBPJ WT rescue identified different Notch-related GO terms (Fig S1B and Table S3), and similarly, a KEGG analysis also identified the “Notch signalling pathway” (adjusted p-value = 0.002575036; mmu04330). We further validated the RNA-Seq data via qPCR focusing on the target genes *Ccdc112, Aig1 and Pmm1* (Fig S1C). To further demonstrate that these genes are regulated by L3MBTL3 in MT cells, we performed L3MBTL3 shRNA knockdown (Fig S2A) and observed that genes, which are significantly downregulated upon rescue with RBPJ WT, but not with RBPJ F261A/A284V mutant, are also upregulated upon L3MBTL3 knockdown (Figs 7E and S2B). Additionally, we performed RNA-Seq analysis of L3MBTL3 depleted MT cells (Figs 7F, S2C-D, and Tables S1, S4, and S5) and observed that a GSEA analysis comparing L3MBTL3-specific shRNA (L3MBTL3 KD) versus control identified for the “Notch signaling pathway” (GO:0007219) a statistically significant and concordant difference between the conditions. The positive NES (normalized enrichment score) indicates an overall induction of Notch target genes (Fig 7F and Table S4) upon loss of L3MBTL3.

## Discussion

Canonical Notch signaling ultimately results in changes in gene expression, which is mediated through the transcription factor CSL^1,2,5^, RBPJ in mammals. RBPJ can function as both a transcriptional repressor or activator by forming structurally similar, but functionally distinct coregulator complexes^7,12^. A recurring theme in RBPJ-coregulator complexes is an ∼15 residue extended peptide, which binds across the front face of the BTD of RBPJ (Fig 1B)^3^. As first shown in NOTCH receptors, the RAM domain forms a high affinity (∼20 nM) interaction with the BTD, which is anchored by the hydrophobic tetrapeptide sequence (-ΦWΦP-, Φ = nonpolar residue) and marks the first step in formation of the Notch ternary activation complex with Mastermind^14^. Corepressors, such as FHL1 and RITA1, also bind to the BTD of RBPJ through RAM-like peptides^18,19^, which contain a -ΦWΦP- motif (Fig 4B), although the affinity of these complexes can vary widely from single-digit micromolar to double-digit nanomolar K_d_’s. The recent structural characterization of the corepressor SHARP revealed a variation of this theme, in which bipartite binding interactions are formed with both the BTD and CTD of RBPJ^20,37^. The avidity of the bipartite interaction leads to an overall affinity of ∼11 nM, and in this way, SHARP sidesteps the strict sequence homology rules for the -ΦWΦP- and other RAM-like regions that bind the BTD^20,37^.

Here we report the 2.06 Å X-ray structure of the RBPJ-L3MBTL3-DNA corepressor complex (Fig 3), which demonstrates that L3MBTL3 interacts with the BTD of RBPJ similar to the RAM domain of NICD, as well as the corepressors FHL1, RITA1, and SHARP, but also displays some unique structural features (Fig 4). L3MBTL3 residue W64 occupies the same conserved position within the hydrophobic tetrapeptide region of all BTD-binders except SHARP, which has a serine at this position (Fig 4B). However, in contrast to other BTD-binders, L3MBTL3 does not have a proline in the fourth position of the hydrophobic tetrapeptide, which is conserved in all BTD-binders, except SHARP, but instead buries a valine sidechain in the corresponding pocket of the BTD that is structurally similar to the isoleucine of SHARP at this position. Several other L3MBTL3 residues bind RBPJ in a structurally similar manner to other coregulators, including P67, which is directly downstream of the hydrophobic tetrapeptide and is conserved in RITA1, SHARP, and some NOTCH orthologs (Fig 4B). Similarly, A60 and T63 of L3MBTL3 structurally align with the corresponding residues from other BTD-binders, following the rules of having a small and branched side chain, respectively, at these positions.

However, unique to L3MBTL3, there are two intervening residues, T61 and T62, which have not been observed in any other RBPJ interacting coregulators (Fig 4B). This insertion forces the L3MBTL3 peptide backbone to bulge outward from the BTD as it passes over a valley between the top and front faces of the BTD (Fig 4C-D). Interestingly, when we determined the structure of the L3MBTL3 Δ62 construct bound to RBPJ, rather than maintaining this bulging conformation, L3MBTL3 Δ62 assumed a conformation more closely resembling other BTD-binders. Importantly, this triple threonine motif is highly conserved in all vertebrate L3MBTL3 orthologs, suggesting it is important for function and could potentially be phosphorylated or O-linked glycosylated; however, to date, there have been no reports of such post translational modification of these residues in L3MBTL3. Taken together, the distinctive structural features L3MBTL3 adopts when in complex with RBPJ greatly expand upon the potential sequences that could bind to RBPJ *in vivo* and may reveal possible modes of regulation via PTMs, which is an understudied area of Notch signaling^38^.

We performed a rigorous ITC binding analysis of the role that each L3MBTL3 residue plays in complex formation with RBPJ, which demonstrated that the determinants of binding are spread across the RBP-ID of L3MBTL3 (Fig 5). Alanine point mutations spanning the L3MBTL3 peptide lead to decreased binding, with several mutants in the extended hydrophobic peptide region (-TWMVP-), except M65, abolishing binding completely, which we corroborated in cells (Fig 6). This is similar to other BTD-binders, demonstrating that the hydrophobic peptide anchors the interaction to RBPJ. The T62A mutation, and to a lesser extent N54A and A58R, surprisingly leads to an increase in binding by the mutant L3MBTL3 peptides. Selective pressure to retain a threonine at position 62 thus creates both the peptide bulge and weakens binding to RBPJ, which may affect the competition of L3MBTL3 with other coregulators, such as NICD upon activation, or alternatively, as mentioned above is functionally important for regulation, *e*.*g*. phosphorylation.

To validate our findings in cells, we used an RBPJ-deficient mature T (MT) cell line^20^ and re-expressed wild-type RBPJ or RBPJ mutants (F261A, A284V, or F261A/A284V) that severely disrupt binding to L3MBTL3 *in vitro* (Table 4 and Fig 7). In this cellular assay, WT RBPJ, but not F261A, A284V, or F261A/A284V, induced repression of the Notch target genes *Lgmn, Hes1*, and *Hey1* (Fig 7B). Consistent with these results, shRNA-mediated knockdown (KD) of L3MBTL3 led to a concomitant de-repression of the targets *Lgmn, Hes1*, and *Hey1* (Fig 7E). Taken together, these studies strongly suggest the involvement of L3MBTL3 in the repression of Notch target genes in cells, albeit we cannot wholly exclude the contributions of other corepressors that bind RBPJ, such SHARP, to the observed RBPJ-mediated repression of target genes.

To further characterize the function of L3MBTL3, we performed RNA-Seq on RBPJ-rescued and L3MBTL3 KD MT cells, which identified the novel targets *Ccdc12, Aig1*, and *Pmm1*. Interestingly, *Ccdc12* has been shown to be involved in erythroid differentiation^39^, which is consistent with the lethality observed in L3MBTL3 knockout mice, whereby a loss in myeloid progenitor differentiation during embryogenesis leads to death by anemia^27^. While this connection remains to be confirmed, this potentially expands our knowledge of L3MBTL3 function.

Finally, Notch transcription complexes are currently being investigated as druggable targets for the treatment of Notch driven diseases^40,41^. However, targeting a conserved binding pocket on the BTD of RBPJ that is used by both coactivators, such as the RAM domain of NICD, and several corepressors, such as L3MBTL3 and SHARP, raises several questions regarding the signaling outcome in cells and the overall utility of this approach^12,38^. Indeed, the recently reported small molecule RIN1 (RBPJ Inhibitor-1) was shown to block the interactions of both NICD and SHARP with RBPJ in cellular assays^40^. These confounding results underscore the importance of a detailed structural and biophysical understanding of how coregulators interact with RBPJ, and how small molecules may tip the balance between repression and activation in different cellular contexts. Certainly, the unusual peptide conformation of L3MBTL3 when bound to RBPJ may lend itself to specific inhibition by targeted small molecules; however, as the first RBPJ targeted small molecules have only recently been identified, further studies are unquestionably required.

## Supporting information

PDB Validation Report

## Data availability

The structures have been deposited in Protein Data Bank (PDB) with accession numbers 7RTE and 7RTI. RNA-Seq data have been deposited at Gene Expression Omnibus (GEO) under the accession number GSE.

## Acknowledgements

We thank members of the Kovall lab for their constructive criticism and the beamline staff at LS-CAT for their technical assistance. We are grateful to P. Käse and T. Schmidt-Wöll for excellent technical assistance.

RAK, DH, BDG and TB designed experiments. DH, BDG, FF performed experiments and analyzed data. DH and ZY performed structural and biophysical studies. BDG, TF and MB performed the bioinformatic analysis. RAK and DH wrote the manuscript, and all authors edited the manuscript.

## Materials and Methods

### Cloning, expression, and protein purification

*Mus musculus* CSL (RBPJ), residues 53-474 (structural core domain), was cloned into both the pGEX-6P-1 vector and a modified pET 28b(+) vector termed pSMT3. The former vector encodes a glutathione S-transferase (GST) fusion protein that can be removed proteolytically with Prescission Protease (GE Healthcare) after affinity purification, leaving the non-native N-terminal sequence GPLGS-. The latter encodes a fusion protein with a His-tagged SMT3 (Suppressor of Mif2 temperature-sensitive mutant 3) construct, which can be cleaved with Ulp1 protease, leaving a single N-terminal serine. Expression and purification were performed as previously described^14,18-20,37^. Transformed bacteria were grown at 37 °C in LB medium, cooled to 10°C, then 2% ethanol and 0.1mM isopropyl β-thiogalactopyranoside (IPTG) were added and induction allowed to occur overnight at 20°C. Bacteria were centrifuged and resuspended in either phosphate buffered saline (PBS) for GST-RBPJ or lysis buffer (20mM Tris pH 8.0, 0.5M NaCl, 50mM Imidazole) for His-SMT3-RBPJ. To purify RBPJ, bacteria were lysed by sonication, the lysate was cleared by centrifugation, and a 3M ammonium sulfate cut precipitated the majority of soluble protein. Resuspended protein was loaded onto glutathione-Sepharose resin or Ni-NTA resin and eluted with either reduced glutathione or imidazole, respectively, then the tags were proteolytically cleaved in manufacturer suggested buffers. Similarly, human L3MBTL3 fragment 1-523, 1-197, and 198-523 were cloned and purified from the pGEX-6P-1 vector. The constructs were further purified to homogeneity using ion exchange and size exclusion chromatography. L3MBTL3 peptides for ITC and crystallography were purchased as HPLC purified synthetic peptides from Peptide 2.0 and received as lyophilized powder.

### Circular dichroism

L3MBTL3 1-197 was dialyzed into a buffer containing 50mM sodium phosphate and 150mM sodium chloride at a concentration of 1.6 mg/ml. Triplicate CD measurements were taken using an Aviv Circular Dichroism Spectrometer Model 215 at 25°C in a 0.01 cm cuvette. The wavelength was scanned from 300 nm to 190 nm in 1 nm increments. CD data were processed using DICHROWEB^42^ and reference set 7 of the CDSSTR^43^ analysis program.

### Isothermal titration calorimetry

ITC experiments were performed at 25°C using a Microcal VP-ITC microcalorimeter. Reaction cell and syringe samples were buffer matched in 50mM sodium phosphate pH 6.5, 150mM NaCl buffer. For the K56A L3MBTL3 mutant, the peptide was dissolved in DMSO first and diluted to 5% DMSO final concentration, while DMSO was added directly to RBPJ for the corresponding experiment. For all binding reactions, 10-15µM RBPJ was used in the cell and 100-150µM L3MBTL3 was used in the syringe. Titrations generally consisted of a single 1µL injection followed by 20-22 14µL injections. The collected data were analyzed using ORIGIN software and fit to a one site binding model.

### Crystallization and data collection

A 15-mer DNA duplex (-TTACCGTGGGAAAGA-/-AATCTTTCCCACGGT-) containing a modified *Hes-1* promoter RBPJ binding site with single-strand TT/AA overhangs was generated by purifying the single stranded oligonucleotides by ion exchange chromatography and annealing in a 1:1 ratio. RBPJ purified from the pSMT3 construct was incubated with the DNA duplex and human L3MBTL3 55-70 wild-type peptide in a 1:1.1:1.1 ratio and screened for crystallization conditions at 4°C using both vapor diffusion and under oil crystallization methods. Vapor diffusion screening was performed in-house on an Art Robbins Phoenix Crystallization Robot. Under oil screening was outsourced to Hauptman Woodward Medical Research Institute’s High Throughput Crystallization Screening Center. Several optimized conditions grew large, diffraction quality crystals with the same space group and resolution despite different morphologies. The reported structure came from a crystallization condition comprised of a 6:10 ratio of Hampton Silver Bullet D3 (0.06 M MES monohydrate, 0.06 M PIPES, 0.33% w/v Hexamminecobalt(III) chloride, 0.02 M HEPES sodium pH 6.8) and 0.1M HEPES pH 6.8, 30% PEG 3350. L3MBTL3 Δ62 crystals were grown in 0.2M ammonium fluoride and 20% PEG 3350. All crystals were cryoprotected with 20% xylitol and flash frozen in liquid nitrogen. Remote data collection occurred at the LS-CAT and NE-CAT beamlines of Advanced Photon Source at Argonne National Lab. Both crystals belong to the orthorhombic P2_1_2_1_2_1_ space group and diffract to under 2.1Å with unit cell dimensions: 67.8Å, 97.1Å, 105.5Å for L3MBTL3 wild-type and 67.9Å, 96.9Å, 105.8Å for L3MBTL3 Δ62.

### Structure determination, model building, and refinement

Diffraction data was processed and scaled using Mosflm^44^ and CCP4i^45^. Phaser^46^ was used to solve the structure via molecular replacement with the RBPJ-DNA complex (PDB: 3IAG)^33^ as a search model. Coot^47^ was used to iteratively build the L3MBTL3 peptide into the model. Refinement was performed using translation/libration/screw parameters in the refine function of Phenix software^48^. Structure validation was performed with Molprobity^49^. The final model contained RBPJ residues 53-473 as well as the residual N-terminal serine. The model also contained L3MBTL3 residues 56-69, leaving off one terminal residue from each end of the peptide. The full DNA duplex is modeled. The structure was refined to R_work_ = 19.9 and R_free_ = 24.3. The PDBePISA server (http://www.ebi.ac.uk/pdbe/pisa/)^34^ was used to calculate the L3MBTL3 binding pocket on RBPJ. Finally, PyMOL (The PyMOL Molecular Graphics System, Version 2.0 Schrödinger, LLC) was used to present structural images and alignments.

### Cell culture, transfection and infection

The mouse hybridoma mature T cell line (MT) was previously described^50,51^ and cultivated in Iscove’s Modified Dulbecco Medium (IMDM, Gibco 21980-065) supplemented with 2 % FCS, 0.3 mg/l peptone, 5 mg/l insulin, nonessential amino acids and penicillin/streptomycin. The MT cells depleted of RBPJ making use of the CRISPR/Cas9 technology were previously described^20^. 293T and Phoenix™ packaging cells (Orbigen, Inc., San Diego, CA, USA) were cultivated in Dulbecco’s modified eagle medium (DMEM, Gibco 61965-059) supplemented with 10% FCS and penicillin/streptomycin. Cells were grown at 37°C with 5% CO_2_. Transfection of Phoenix™ cells and retroviral infection of MT cells were performed as previously described^52^. Transfection of 293T cells, lentiviral infection and selection of MT cells were performed as previously described^53^. U87-MG cells were cultivated in DMEM medium supplemented with 10% FBS and penicillin/streptomycin as previously described^21^.

### Constructs

The pcDNA3.1 Flag-mRBPJ WT CRISPR/Cas9 resistant (CRr), the pMY-Bio IRES Blasticidin, the pMY-Bio-Flag-mRBPJ WT CRr IRES Blasticidin and the pMY-Bio-Flag-mRBPJ F261A CRr IRES Blasticidin were previously described^20^. The pcDNA3.1 Flag-mRBPJ A284V CRr and the pcDNA3.1 Flag-mRBPJ F261A/A284V CRr were generated via site directed mutagenesis using the QuikChange II XL Site-Directed Mutagenesis Kit (Agilent Technologies 200521-5) accordingly to manufacturer’s instructions with the oligos listed in Table S6 and using the pcDNA3.1 Flag-mRBPJ WT CRr and the pcDNA3.1 Flag-mRBPJ A284V CRr as templates, respectively. The pMY-Bio-Flag-mRBPJ A284V CRr IRES Blasticidin and the pMY-Bio-Flag-mRBPJ F261A/A284V CRr IRES Blasticidin were generated via restriction digestion. Briefly, the pcDNA3.1 Flag-mRBPJ A284V CRr and the pcDNA3.1 Flag-mRBPJ F261A/A284V CRr were digested with NotI (NEB) and the cDNAs were inserted into the pMY-Bio IRES Blasticidin pre-digested with NotI (NEB). The reporter construct 5 x Gal4-RE-LUC (pFR-Luc) was described previously^36^. The Gal4 expression vector pFA-CMV (Agilent/Stratagene) was used as control and as cloning vector for the Gal4-L3MBTL3 fusions. PCR fragments were digested with *EcoRI* and *HinDIII* and inserted into the corresponding sites of pFa-CMV, resulting in the Gal4-L3MBTL3 (1-197) fusion constructs (wt, T63A, W64A, M65A, V66A, P67A and 5xA). The pcDNA3-HA-tagged-L3MBTL3 expression vector was described previously^21^ and site-directed mutagenesis was used to create L3MBTL3 mutants. All oligonucleotides used in this study are listed in Table S6. All plasmids were analyzed by sanger sequencing.

### ShRNA knockdown

For the knockdown in MT cells, the pLKO.1 TRC1 shRNA library (SIGMA-ALDRICH) was used. Sequence of the hairpin is indicated in Table S6.

### RNA extraction, RT-PCR and qPCR from cell lines

Total RNA was purified using Trizol reagent (Ambion 15596018) accordingly to manufacturer’s instructions and 1 μg of RNA was reverse-transcribed into cDNA using M-MuLV reverse transcriptase (New England Biolabs) and random hexamers. qPCRs were performed using gene-specific oligonucleotides, double-dye probes (see Table S6), Absolute QPCR ROX Mix (Thermo Scientific AB-1139), and analyzed using the StepOnePlus™ Real-Time PCR System (Applied Biosystem). Data were normalized to the housekeeping gene *Glucuronidase* β (GusB). Alternatively, RNA was purified using the RNeasy Mini Kit (Qiagen #74104), the QIAshredder (Qiagen #79654) and treatment with DNase I (Qiagen #79254) accordingly to manufacturer’
ss instructions.

### Protein extract, CoIP, cell fractionation and Western blotting

Whole Cell Extract (WCE) from MT cells was prepared as follows. Briefly, cells were washed twice in PBS, lysed in WCE buffer (20 mM Tris-HCl pH 8.0, 150 mM NaCl, 1 % NP-40, 10 % glycerol, 0.5 mM Na_3_VO_4_, 10 mM NaF, 1 mM PMSF, 1x protease inhibitor cocktail mix) and incubated 20 min on ice. Samples were centrifuged 15 min at 13200 rpm at 4°C and protein concentration measured by Bradford assay (Sigma-Aldrich).

The nuclear extract of MT cells was prepared as follows. Briefly, cells were washed twice with PBS, resuspended in Hypotonic buffer (20 mM Hepes, 20 mM NaCl, 10 % glycerol, 5 mM MgCl_2_, 0.2 mM PMSF) and incubated 20 min on ice. Cell suspensions were vortexed and lysates were centrifuged at 4000 rpm 10 min at 4°C. After collecting the supernatant (cytoplasm), the pellets (nuclei) were washed with PBS and lysed in Hypertonic buffer 300 mM NaCl (20 mM Hepes pH 7.9, 300 mM NaCl, 0.3 % NP-40, 25 % glycerol, 1 mM MgCl_2_, 0.2 mM PMSF, 1x protease inhibitor cocktail mix, 0.3 mM DTT). Samples were incubated 20 min on ice in cold room, centrifuged at 13000 rpm 5 min at 4°C and protein concentration was measured by Bradford assay. Proteins were separated by SDS-PAGE and transferred to a nitrocellulose membrane (Amersham 10600006) using the Biorad Mini Trans-Blot system. The RBPJ (Cosmo Bio Co., Clone T6709) and TBP (Santa Cruz sc-273) Western blotting were performed essentially as previously described^54^. In the case of the L3MBTL3 (Bethyl A302-852A) Western blotting, membranes were incubated 1 h at room temperature in blocking solution (5 % nonfat dry milk, 1x TBS, 0.1 % Tween 20) and incubated over night with primary antibody diluted 1:5000 in blocking solution. Membranes were washed five times in 1x TBS, 0.1 % Tween 20 and incubated 1 h at room temperature with the proper secondary antibody diluted 1:5000 in blocking solution. Membranes were washed five times in 1x TBS, 0.1 % Tween 20. All membranes were incubated at room temperature with ECL solution and signals were acquired with a Vilber Fusion FX7 system. The following secondary antibody were used: anti-rat IgG HRP (Jackson ImmunoResearch, 112-035-072) and anti-rabbit IgG HRP (Cell Signaling #7074S).

### Mammalian Two Hybrid Luciferase assay

HeLa cells were seeded in 48-well plates at a density of 20 × 10^4^ cells. Transfection of the reporter construct pFR-Luc (5 × Gal4-RE-LUC) together with Gal4 or Gal4-L3MBTL3 expression constructs was performed with Lipofectamine 2000 reagent (Thermo Fisher Scientific) using 250 ng of reporter plasmid alone or together with increasing amounts of expression plasmid (50 ng, 100 ng). After 24 hours luciferase activity was determined from at least four independent experiments from 20 μl of cleared lysate. Measurements were performed using a LB 9501 luminometer (Berthold) and the luciferase assay system from Promega.

### RNA-seq data analysis

The systemPipeR R/BioC package with customized parameter files was used to generate system calls within R^55^. Raw sequencing reads were aligned to the mouse genome (mm9) and the corresponding GTF file (downloaded from Illumina’s IGenomes site) using TopHat v.2.1.1 with parameters i= 30, I = 3000 and g = 1 and alignments were stored as BAM files^54^. These BAM files and the gene annotation were used to calculate the gene-specific count tables for all samples with the summarizeOverlaps function^56^. The normalization (including batch effects) of resulting count tables per gene and subsequent detection of deregulated genes was done using DESeq2 v.1.24.0 with default settings^57^. Pearson correlation coefficient (PCC) was calculated based on the significantly deregulated genes (for RBPJ rescue: False discovery rate < 0.05 and log2FoldChange < -0.5 or > 0.5; for L3MBTL3 KD: False discovery rate < 0.05 and log2FoldChange < - 1 or > 1). RBPJ target genes were chosen as those genes, which were significantly downregulated upon rescue with RBPJ WT compared to the eV control. GO, GSEA, and KEGG analysis were done within R using clusterProfiler^58^ with standard parameters and adjusted p-values cutoffs of 0.1 (L3MBTL3 KD) or 0.01 (L3MBTL3 rescue). Genes were ranked based on the Wald test statistic resulting from DESeq2 analysis (see above). The universe for this analysis was defined as all genes that have detectable read counts in at least one sample. Analysis code is available upon request.

