## Supplemental Data for "The structure, binding, and function of a Notch transcription complex involving RBPJ and the epigenetic reader protein L3MBTL3"

### Supplemental Information

**Table S1. Summary statistics for RNA-Seq experiments performed in this study.** Table showing the TopHat alignment statistics for RNA-Seq experiments including number of reads (Nreads), number of aligned reads (Nalign) and the percentage of alignment (Perc_Aligned) per individual sample.

**Table S2. RNA-Seq analysis of mature T (MT) cells depleted of endogenous RBPJ infected with empty vector (eV) as a control, RBPJ WT or RBPJ F261A/A284V double mutant.** Table of the DESeq2 statistics including three contrasts (RBPJ WT / eV, RBPJ F261A/A284V / eV and RBPJ F261A/A284V / RBPJ WT) and the normalized read counts for all samples per gene.

**Table S3. Gene Ontology (GO) analysis (Biological Pathways) of significant downregulated genes upon RBPJ WT rescue in mature T (MT) cells depleted of endogenous RBPJ.** Table showing the results of the gene ontology (GO) analysis including biological and statistical information (GO ID, Description, GeneRatio, BgRatio, pvalue, p.adjust, qvalue, geneID and Counts).

**Table S4. RNA-Seq analysis of mature T (MT) cells infected with hairpin directed against L3MBTL3 or scramble (Control) as a control.** Table of the DESeq2 statistics including the contrast (L3MBTL3 KD / Control) and the normalized read counts for all samples per gene.

**Table S5. GSEA analysis of differentially upregulated genes upon L3MBTL3 depletion in mature T (MT) cells.** Table of GSEA analysis (GO biological process) results of the L3MBTL3 KD / control RNA-Seq experiment.

**Table S6.** Oligonucleotides used for cloning purposes, shRNAs and qPCR experiments.

| | | |
|---|---|---|
| **shRNA** | | |
| L3MBTL3 KD | 5´-CCG CTC ACT GAG CTA AAC TTA-3´ | |
| **Site directed mutagenesis** | | |
| RBPJ A284V fw | 5´-GTG CTC AGT GAC TGG CAT GGT ACT CCC AAG ATT GAT AAT TAG-3´ | |
| RBPJ A284V rev | 5´-CTA ATT ATC AAT CTT GGG AGT ACC ATG CCA GTC ACT GAG CAC-3´ | |
| mRBPJ F261A fw | 5´-GAA AGT GAG GGG GAA GAG GCC ACA GTT AGA GAT GGC TAC-3´ | |
| mRBPJ F261A rv | 5´-GTA GCC ATC TCT AAC TGT GGC CTC TTC CCC CTC ACT TTC-3´ | |
| **Gal4-L3MBTL3 fusion constructs** | | |
| Gal4-L3MBTL3_fwd | 5´-ATG AAT TCT AAC GGA ATC TGC CTC TAG CAC AAG-3´; | |
| Gal4-L3MBTL3_rev | 5´-ATA AGC TTT TAC TGC TTC AGT ACA GCC GAA TCC-3 | |
| **real time PCR** | | |
| | *Mus musculus* | *Probe* |
| *Aig1* | 5'-AAT GCC CTC GCA TCA GAC-3' | #16 |
| | 5'-AAG AAC ACG GCC TGG ATA AC-3' | |
| *Ccdc112* | 5'-ACC AAA GAG GCG ACT TCA GA-3' | #4 |
| | 5'-CTG TTG CTG GAT TTT TGC TCT-3' | |
| *GusB* | 5´-TGT GGG CAT TGT GCT ACC T-3´ | #25 |
| | 5´-ATT TTT GTC CCG GCG AAC-3´ | |
| *Hes1* | 5´-TGC CAG CTG ATA TAA TGG AGA A-3´ | #20 |
| | 5´-CCA TGA TAG GCT TTG ATG ACT TT-3´ | |
| *Hey1* | 5´-CAT GAA GAG AGC TCA CCC AGA-3´ | #17 |
| | 5´-CGC CGA ACT CAA GTT TCC-3´ | |
| *Lgmn* | 5´-GAA TTC CCA CGG TTC TGC-3´ | #85 |
| | 5´-AGC ACC AGG CTG AGA AGC-3´ | |
| *Pmm1* | 5'-GGT ACA GAT CGG CGT GGT-3' | #21 |
| | 5'-ACA CGT AGT CAA ACT TCT CAA TGA CT-3' | |
| *Tbp* | 5´-GGG GAG CTG TGA TGT GAA GT-3´ | #97 |
| | 5´-CCA GGA AAT AAT TCT GGC TCA T-3´ | |

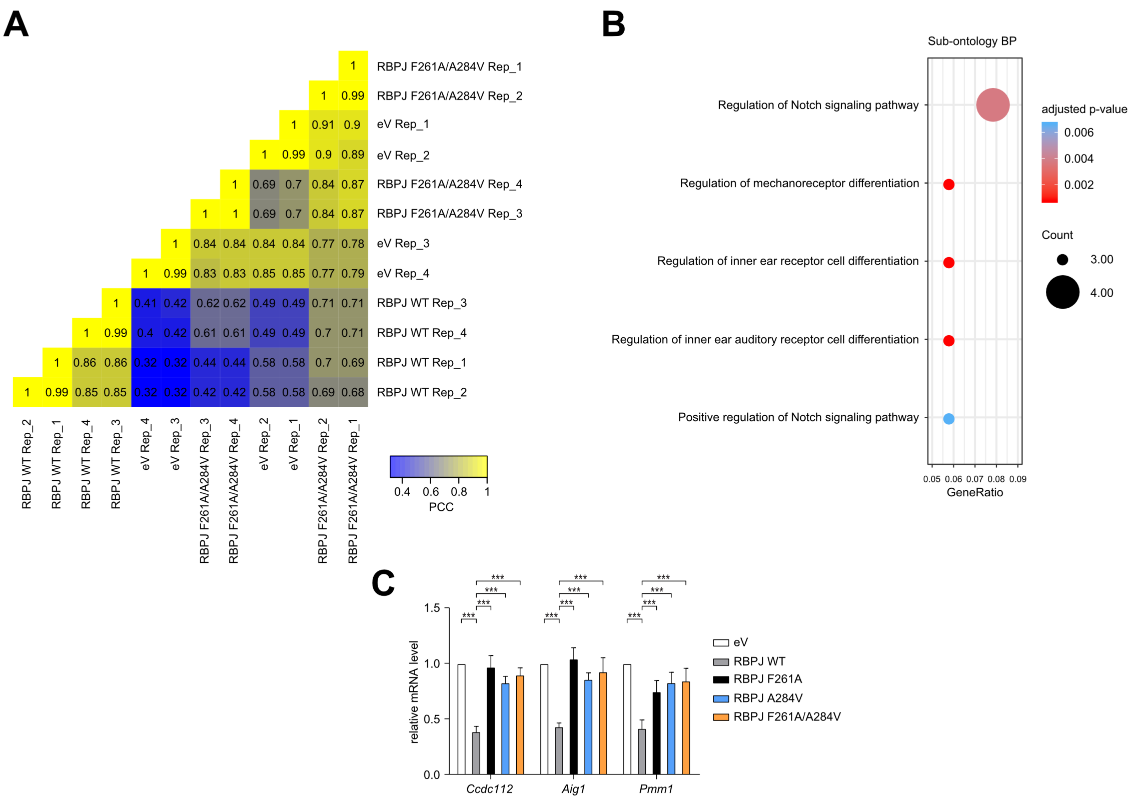

**Figure S1. RBPJ function can be restored via overexpression of RBPJ WT but not of F261A/A284V mutant in mature T (MT) cells depleted of endogenous RBPJ.** (A) Heat map of the Pearson correlation coefficients (PCC) based on pair-wise comparisons of the normalized read counts of all significantly deregulated genes (FDR $< 0.05$ and log2FoldChange-value $< -0.5$ or $> 0.5$ when comparing RBPJ rescue with empty vector control). (B) Gene ontology (GO) analysis (sub-ontology "Biological Process") of the significant downregulated genes upon RBPJ WT rescue in mature T (MT) cells depleted of endogenous RBPJ identified Notch-related GO terms. (C) Expression of Notch target genes is downregulated by RBPJ WT but not by the single or double mutants in MT cells depleted of endogenous RBPJ. MT cells depleted of RBPJ were infected with viruses carrying plasmids encoding for RBPJ WT, RBPJ F261A, RBPJ A284V, RBPJ F261A/A284V or empty vector (eV) as control. Upon RNA extraction and reverse transcription, cDNAs were analyzed by qPCR using assays specific for *Ccdc112*, *Aig1* or *Pmm1*. Data were normalized versus the housekeeping gene *GusB*. Shown is the mean ± SD of seven experiments (***$p < 0.001$, unpaired Student's t test).

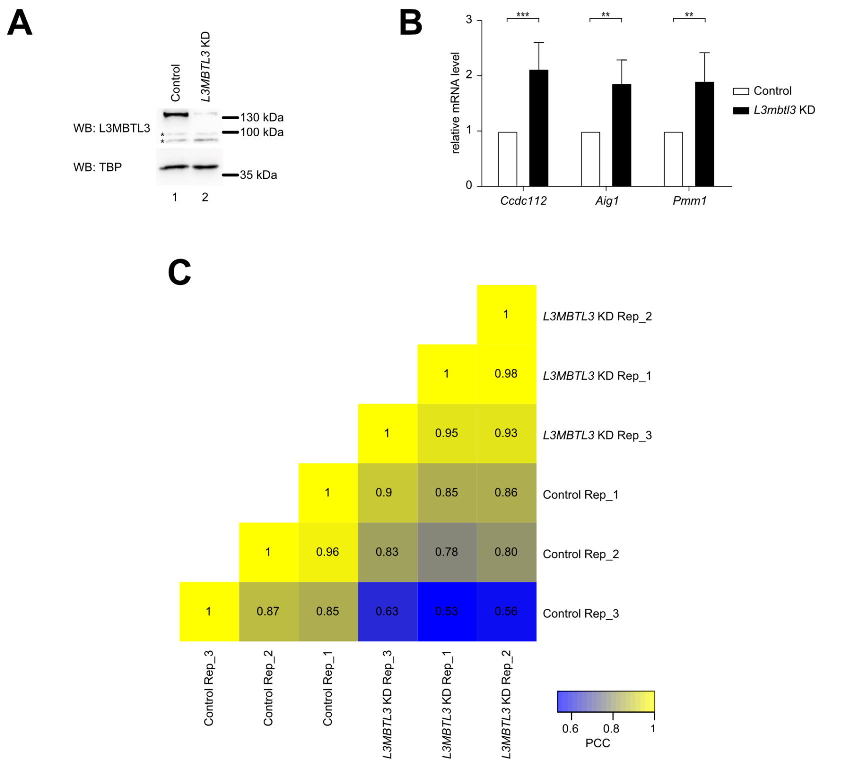

**Figure S2. L3MBTL3 is a positive regulator of the Notch signaling pathway.** (A) L3MBTL3 depletion in mature (MT) cells leads to upregulation of Notch target genes. MT cells were infected with short hairpin RNAs directed against L3MBTL3 or scramble (Control) as a control. Whole cell extracts (WCE) were analysed by Western blotting (WB) with the indicated antibody. TBP was used as a loading control. (B). L3MBTL3 depletion in MT cells leads to upregulation of Notch target genes. MT cells were infected with hairpin directed against L3MBTL3 or scramble (Control) as a control. Upon RNA extraction and reverse transcription, cDNAs were analysed by qPCR using assays specific for *Ccdc112*, *Aig1* or *Pmm1*. Data were normalized versus the housekeeping gene *GusB*. Shown is the mean ± SD of five experiments (**p < 0.01, ***p < 0.001, unpaired Student's t test). (C) Heat map of the Pearson correlation coefficient (PCC) for pair-wise comparison of the normalized read counts of all significantly deregulated genes (FDR < 0.05 and log2FoldChange-value < -1 or > 1 when comparing L3MBTL3 knock down with control).
